# Phase-separated FRM proteins create signaling subdomains in inter-organelle junctions

**DOI:** 10.64898/2026.11.24.769035

**Authors:** Huichao Deng, Jinbo Cheng, Richard D. Fetter, Xiaoyu Tang, Alex Calhoun, Wendy Herbst, Guanghui Qin, Xinyue Li, Can Wang, Raymond Dunn, Jianxiu Zhang, Xing Liang, Caitlin A Taylor, Jessica Feldman, Saul Kato, Mingjie Zhang, Hang Lu, Meijing Li, Xiandeng Wu, Kang Shen

## Abstract

Endoplasmic reticulum-plasma membrane junctions (EPJs) house the calcium-signaling machinery that mediates Ca²⁺ release from the ER and its reentry into the ER. How these opposing functions are coordinated at EPJs is unknown. Here we show that neuronal EPJs are patch-like membrane contacts partitioned into discrete, adjacent subdomains containing the Ca²⁺-release and Ca²⁺-reentry machineries. Through a forward genetic screen, we identified the evolutionarily conserved FERM-family proteins FRM-4 and FRM-1 as central organizers of this architecture. FRM-4 and FRM-1 assemble EPJs by bridging the membranes, recruiting Ca²⁺-signaling components, and establishing the subdomains. Loss of FRM proteins disrupts both signaling domains and attenuates neuronal Ca²⁺ responses and plasticity. FRM proteins and Ca²⁺-signaling molecules undergo phase separation at EPJs *in vivo*. Purified FRM proteins phase separate *in vitro* and suffice to partition Ca²⁺ channels into their correct subdomains. Thus, phase separation creates molecular and functional subdomains at organelle junctions.

**Highlights:** ▪ Somatic neuronal ER-PM junctions form a patchwork of discrete membrane contacts

▪ SOCE and CICR components occupy adjacent, molecularly distinct subdomains

▪ FRM-4 and FRM-1 organize junction architecture and calcium-signaling proteins

▪ FRM condensates selectively partition signaling proteins and support Ca^2+^ responses

## Introduction

Endoplasmic reticulum-plasma membrane junctions (EPJs) are membrane contact sites where endoplasmic reticulum (ER) is closely juxtaposed against the plasma membrane (PM).^1–4^ Widely conserved across eukaryotes,^5–7^ EPJs support non-vesicular lipid transfer,^8–16^ ion-channel clustering^17–19^ and prominently, calcium signaling by placing proteins on opposing membranes within nanometers of one another.^18,20–24^ In neurons, EPJs cover a substantial fraction of the somatic surface, localizing along dendrites at ∼1 µm intervals, and are present in presynaptic boutons.^3,25,26^ Their composition, however, has been inferred largely with one tagged protein at a time, and how the many EPJ components are arranged relative to one another within a single neuronal junction is unknown.

Two calcium-signaling pathways use the junctional cleft in opposite directions. Store-operated calcium entry (SOCE) refills a depleted ER store: the ER Ca²⁺ sensors STIM1 and STIM2 detect the fall in luminal Ca²⁺, oligomerize and translocate to EPJs, where they bind and open the plasma membrane Ca²⁺ channel ORAI.^18,22–24,27–37^ Ca²⁺ that enters through ORAI is returned to the ER by SERCA pumps positioned near the junction, forming a local feedback circuit for store homeostasis.^38,39^ STIM proteins also operate in neurons, where they maintain spine Ca²⁺ levels,^40^ shape Purkinje cell signaling^41^ and set presynaptic release probability.^25^ Notably, junctional STIM1 not only activates ORAI but also binds and inhibits L-type voltage-gated Ca²⁺ channels (Ca_V_1).^42,43^

Calcium-induced calcium release (CICR), by contrast, releases Ca²⁺ from a loaded store. Depolarization opens Ca_V_1, and the resulting Ca²⁺ microdomain activates ryanodine receptors (RyRs) in the apposed ER membrane.^19,44–46^ Junctophilins organize this coupling by anchoring in the ER, binding plasma membrane lipids through MORN motifs, and binding Ca_V_1 and RyR directly.^21,47^ The brain isoforms JPH3 and JPH4 assemble Ca_V_1-RyR-K⁺ channel complexes that control hyperpolarization, excitability and learning,^48,49^ and a JPH3 repeat expansion causes Huntington disease-like 2.^50^ In mammalian somata, K_V_2.1-induced EPJs couple Ca_V_1.2 to RyR and downstream PKA and transcriptional signaling,^19,51,52^ and in *Caenorhabditis elegans* (*C. elegans*) the sole junctophilin JPH-1 maintains ER-PM contacts and supports neurotransmission.^53^

SOCE and CICR are thus two halves of one problem - maintaining and deploying the ER Ca²⁺ store, and are built from overlapping parts: both require nanometer apposition, both are organized by lipid-binding tethers, and junctophilins can recruit STIM1 just as STIM1 can bind Ca_V_1.^43,54^ Yet they move Ca²⁺ in opposite directions across the ER membrane, and are mutually antagonistic through STIM1-mediated Ca_V_1 inhibition. Indiscriminate mixing of the two machineries within a shared cleft would risk futile cycling of released and re-sequestered Ca²⁺, or Ca_V_1 inhibition during store refilling. How neurons position SOCE and CICR components relative to one another has not been examined; membrane contact sites have generally been treated as uniform platforms rather than internally patterned structures.

Liquid-liquid phase separation (LLPS) driven by multivalent interactions concentrates specific proteins into compartments without a delimiting membrane,^55–58^ and proceeds at lower threshold concentrations when one component is membrane anchored.^55^ Most mechanistic understanding of condensate specificity comes from reconstitution experiments. *In vitro*, high-valency scaffolds define a phase and lower-valency clients partition according to the sites they can engage, so that small changes in stoichiometry switch a client from included to excluded;^57,59^ in condensates built from folded domains, point mutations at a single interface redirect partitioning.^60,61^ Reconstituted condensates can also exclude, and can coexist as immiscible phases: on supported bilayers, LAT condensates concentrate the kinase ZAP-70 while excluding the phosphatase CD45, segregating opposing enzymatic activities;^58^ *in vitro*, synapsin and RIM/RIM-BP form separate phases that organize reserve versus docked vesicle pools;^62^ and reconstituted CaMKII-GluN2B condensates segregate AMPA receptor PSD-95 complexes into a distinct phase, a behavior recapitulated in heterologous cells and cultured neurons.^63^ Functionally opposite excitatory and inhibitory postsynaptic densities are naturally demixed condensates in dually innervated dendritic spines with sub-micrometer in width.^64^ In the nucleus, HP1α condensates that compact chromatin and exclude activators were characterized *in vitro* and in cells,^65^ with heterochromatin domains in *Drosophila* embryos showing liquid-like behavior *in vivo*.^66^ Collectively these studies suggest that phase separation can sort related but opposing machineries into separate territories, but the sorting itself has been demonstrated almost entirely with purified or overexpressed components.

Whether LLPS performs such functions in living tissue is harder to establish, because direct observation of phase condensate behavior is technically difficult, and condensate-like appearance can arise without phase separation.^67,68^ One of the earliest and most compelling demonstrations of liquid-like condensate behavior *in vivo* came from *C. elegans* embryos, where P granules were shown to undergo fusion, deformation, and spatially regulated dissolution and condensation;^69^ SYD-2/Liprin-α and ELKS-1 form dynamic condensates during early presynaptic active-zone assembly. Mutations that selectively impair phase separation while preserving synaptic localization and tested protein interactions disrupt active-zone assembly and synaptic function in intact animals;^70^ MEC-2 condensates undergo a liquid-to-solid transition required for touch sensation.^71^ These studies established that condensates form and are required *in vivo*, but none has asked whether a condensate sorts distinct signaling machineries into adjacent subdomains in an intact animal.

Here, we combine serial-section transmission electron microscopy (ssTEM), endogenous protein labeling, forward genetics, cryo-electron tomography (cryo-ET), protein reconstitution and calcium imaging in the *C. elegans* PVD sensory neuron to address these questions. We find that somatic EPJs form a patchwork of discrete contacts containing adjacent SOCE- and CICR-associated subdomains. A forward genetic screen identifies two FERM-domain proteins, FRM-4 and FRM-1, as cooperative organizers of these contacts. Endogenously tagged FRM proteins and calcium-signaling components form dynamic condensates at EPJs *in vivo*; purified FRM proteins undergo LLPS *in vitro* and are sufficient to partition SOCE and CICR components into distinct phases. Loss of FRM proteins leads to loss of patch-like EPJs, reduced junctional proteins and impaired stimulus-evoked calcium responses.

## Results

### Neuronal somatic EPJs form a distinctive soccer ball-like architecture

To characterize the ultrastructural organization of neuronal EPJs, we performed ssTEM and three-dimensional (3D) reconstruction of the *C. elegans* PVD neuronal soma (Figures 1A-1D). Unlike the conventional stacked ER cisternae, the somatic ER formed a continuous, barrel-like network beneath the PM (Figure 1A). The ER membrane was extensively decorated with ribosomes, characteristic of rough ER (Figures 1A and 1A.1). Notably, portions of the ER closely apposed against the PM, forming extensive membrane contact sites with two distinguishing ultrastructural features (Figures 1A and 1A.2). First, ribosomes were excluded from the ER membrane facing the PM at these sites (Figures 1A and 1A.2). Second, the distance between ER and PM in the ribosome-free zone is nearly constant, averaging at 23 nm, consistent with the dimensions of previously characterized EPJs.^3,4^ On the contrary, at the edge of the ribosome-free zone, the distance between ER and PM increases to above 50 nm with the appearance of the first ribosome (Figure 1C).

**Figure 1.**
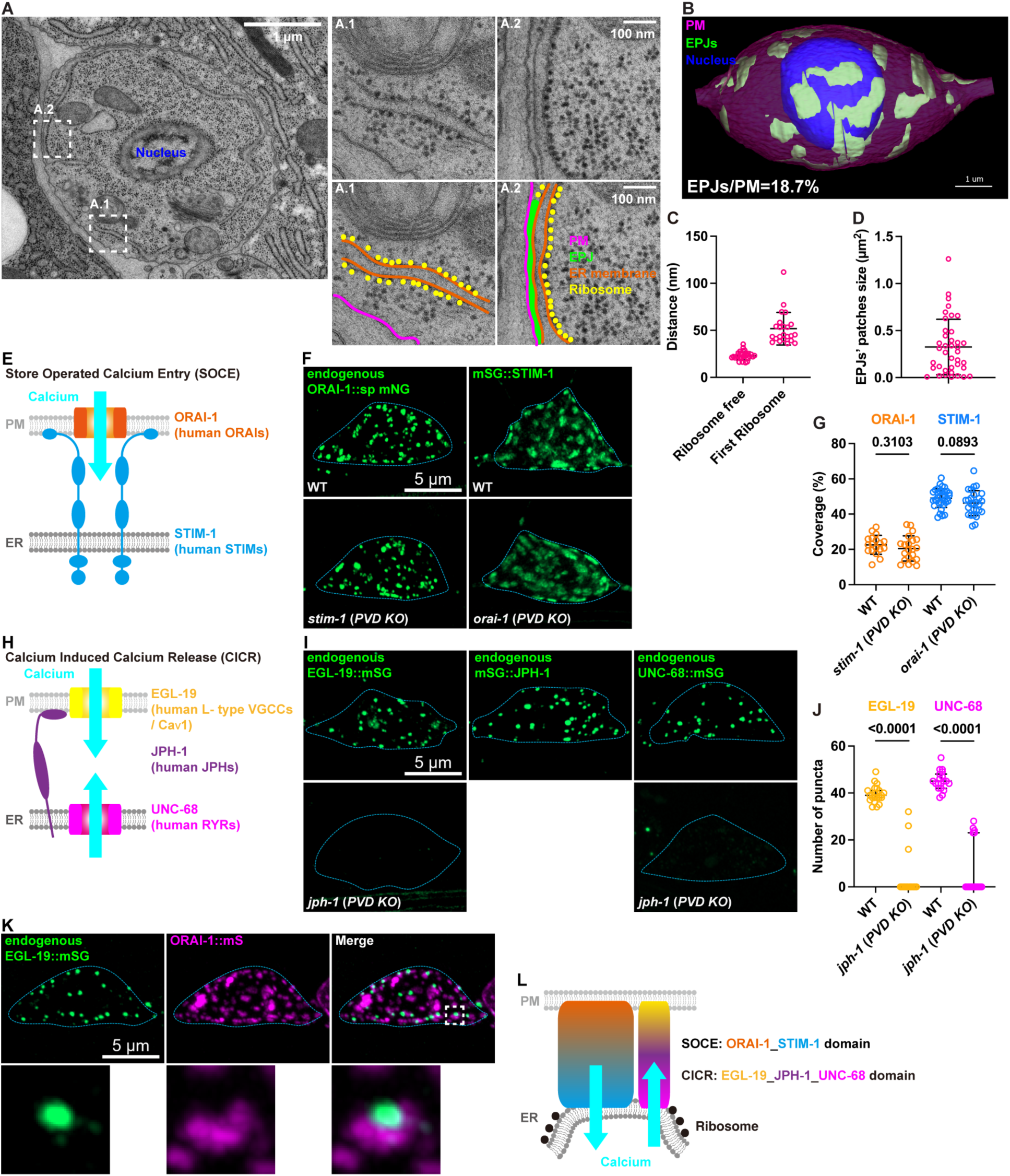
Neuronal EPJs contain adjacent SOCE and CICR subdomains. (A) Representative ssTEM image of an L4-stage *C. elegans* PVD neuronal soma showing cortical rough endoplasmic reticulum (ER) and EPJs. Enlarged regions show ribosome-decorated rough ER (A.1) and an EPJ with close ER-PM apposition and ribosome exclusion from the apposed ER membrane (A.2). (B) 3D reconstruction from ssTEM showing the patch-like distribution of EPJs across the PVD soma. (C) Quantification of the distances from the plasma membrane to ribosome-free ER regions and to the first ribosome on the ER surface in a single reconstructed PVD soma. Each dot represents an individual measurement (ribosome-free regions, n = 30; first ribosomes, n = 23). Mean ± SD: 23.05 ± 4.43 nm and 51.79 ± 17.32 nm, respectively. All measurements were obtained from one PVD soma ssTEM 3D reconstruction. No inferential statistical test was performed. (D) Quantification of individual EPJ patch areas (n = 39) in a representative ssTEM-based 3D reconstruction of the PVD soma. Each dot represents one EPJ patch. Data are presented as mean ± SD. All patches were measured from a single PVD neuron in one animal. (E) Schematic of SOCE components, including the ER Ca²⁺ sensor STIM-1 and PM Ca²⁺ channel ORAI-1. (F) Representative confocal z-stack images of endogenously split-mNeonGreen-labeled ORAI-1 and single copy mStayGold::STIM-1 transgene in PVD somas of the indicated genotypes. (G) Quantification of ORAI-1 and STIM-1 coverage in the PVD soma of WT animals and animals with PVD-specific knockout of *stim-1* or *orai-1*, respectively (ORAI-1: n = 18 and 21; STIM-1: n = 33 and 27, respectively). Each dot represents one PVD neuron from an independent animal. Data are presented as mean ± SD. Statistical significance was determined using two-tailed unpaired Welch’s t tests. (H) Schematic of CICR components: the L-type voltage-gated Ca²⁺ channel EGL-19, junctional protein JPH-1, and ER ryanodine receptor UNC-68. (I) Representative confocal z-stack images of endogenously mStayGold-tagged EGL-19, JPH-1, and UNC-68 in PVD somas of the indicated genotypes. (J) Quantification of endogenous EGL-19 and UNC-68 puncta in PVD neuronal somas of wild-type (WT) and PVD-specific *jph-1* knockout (*PVD KO*) animals. Each dot represents one PVD neuron from an independent animal. EGL-19: WT, n = 23; *jph-1* (*PVD KO*), n = 15. UNC-68: WT, n = 19; *jph-1* (*PVD KO*), n = 18. Data are presented as median with interquartile range (IQR). Statistical significance was determined using two-sided, unpaired Mann-Whitney U tests without adjustment for multiple comparisons. (K) Representative confocal z-stack images showing adjacent distributions of endogenously mStayGold-tagged EGL-19 and ORAI-1::mScarlet expressed from a single-copy transgene in the PVD soma. (L) Model of adjacent, molecularly distinct SOCE and CICR subdomains at neuronal EPJs.

3D reconstruction further revealed that these junctions were not distributed uniformly across the neuronal surface. Instead, they formed discrete, patch-like domains that collectively produced a striking soccer ball-like pattern across the soma (Figures 1B and 1D). These junctional patches occupied approximately 18.7% of the total PM surface area. Similar patchwork architectures were observed in ssTEM reconstructions of additional neuronal somas in the head ganglia (Figure S1A), suggesting that this organization is not unique to PVD neurons. Thus, neuronal somatic EPJs exhibit a distinctive, spatially patterned architecture rather than forming a continuous membrane interface.

### SOCE and CICR machineries segregate into spatially distinct junctional subdomains

The striking patchwork organization of neuronal EPJs prompted us to investigate how Ca²⁺-signaling machinery is distributed within these junctions. At EPJs, SOCE, mediated by the ER Ca²⁺ sensor STIM-1 and the PM Ca²⁺ channel ORAI-1, recharge ER Ca²⁺ (Figure 1E).^72,73^ In parallel, CICR, involving the L-type voltage-gated Ca²⁺ channel EGL-19/ Ca_V_1, the junctional protein JPH-1/ Junctophilin, and the ER Ca²⁺-release channel UNC-68/ RyR, release Ca²⁺ from ER (Figure 1H).^74–76^

To visualize endogenous SOCE components, we inserted split mNeonGreen^77,78^ into the chromosomal locus of *orai-1* using CRISPR,^79^ and mStayGold::STIM-1 expressed from a single copy transgene expression system.^80^ Both proteins formed discrete patch-like structures on Z-projected confocal image stacks (Figure. 1F top row) and punctate patterns on cell cortex in single confocal slices (Figures S2A-S2C). This is reminiscent of the junctional patches revealed by ssTEM (Figure 1B). Surprisingly, disruption of *stim-1* by PVD conditional knock out^81,82^ did not abolish the punctate localization of ORAI-1, nor did loss of *orai-1* affect the junctional distribution of STIM-1 (Figures 1F and 1G). These findings indicate that ORAI-1 and STIM-1 can maintain their spatial organization independently of one another in neuronal soma, contrasting with the canonical STIM-dependent recruitment of ORAI channels during SOCE activation in other cell types.^18,83^

We next examined the spatial organization of CICR machinery by endogenously labeling EGL-19/ Ca_V_1, JPH-1/ Junctophilin, and UNC-68/ RyR with mStayGold. All three proteins formed discrete puncta along cell cortex in single confocal slices (Figures S2D-S2H), and appear as roughly evenly distributed puncta on Z-projected confocal image stacks (Figure 1I top row). The junctional localization of EGL-19/ Ca_V_1 and UNC-68/RyR completely depended on JPH-1 (Figures 1I bottom row and 1J), consistent with the established role of junctophilins in organizing excitation-contraction coupling machinery in muscle cells.^21,47,84^

To determine how these two Ca²⁺-signaling systems are spatially organized relative to one another, we simultaneously visualized endogenously labeled EGL-19::mStayGold and ORAI-1::mScarlet expressed from a single copy transgene expression system. Remarkably, EGL-19 and ORAI-1 signals are closely juxtaposed but mutually exclusive with each other. Thus, instead of a homogeneous signaling platform, neuronal EPJs are partitioned into neighboring molecular subdomains enriched in CICR and SOCE machinery, respectively (Figures 1K and 1L). This spatial segregation raises the question of how these specialized subdomains are established and maintained.

### FRM-4 and FRM-1 cooperatively organize neuronal EPJs

To identify molecular regulators of neuronal EPJ architecture, we expressed the artificial ER-PM tether GFP::MAPPER^9^ in PVD neurons to visualize junctional morphology *in vivo* (Figure S3A). GFP::MAPPER formed discrete patches along the PVD soma periphery, resembling the EPJ distribution revealed by ssTEM (Figure S3A). Using this reporter, we performed a forward genetic screen and isolated the mutant allele *wy1931*, which exhibited markedly fewer and enlarged EPJ patches (Figures S3A-3C). SNP mapping and whole-genome sequencing identified a premature stop codon in the *frm-4* (*Q158X*) gene. To establish causality, we generated a complete deletion of *frm-4* using CRISPR-Cas9. Both *frm-4* (*wy1931*) and *frm-4* (*null*) mutants displayed similar defects in EPJ morphology, characterized by a reduction in junctional patch number (Figures S3A-S3C). Re-expression of wild-type FRM-4 specifically in PVD neurons rescued these defects, demonstrating that FRM-4 acts cell-autonomously to regulate neuronal EPJ architecture (Figures S3A-S3C).

FRM-4 belongs to a family of proteins containing a conserved FERM (4.1 protein, ezrin, radixin, moesin) domain, which is commonly links membrane associated proteins to the cortical actin cytoskeleton.^85,86^ Three FRM-4 paralogs are encoded in the *C. elegans* genome: FRM-1, FRM-2, and ERM-1. Unlike FRM-4, which contains a predicted C-terminal transmembrane (TM) domain, these proteins lack predicted membrane-spanning regions (Figure S3B). To determine whether they also contribute to EPJ organization, we generated complete deletion alleles of *frm-1* and *frm-2* and a PVD-specific knockout of the essential gene *erm-1*. Among these mutants, loss of *frm-1* caused a reduction in EPJ patch number, although the phenotype was less severe than that of *frm-4* (*null*) (Figures S3A-S3C). Notably, *frm-4* (*null*); *frm-1* (*null*) double mutants exhibited a more pronounced reduction in junctional patch number than *frm-4* (*null*) single mutants (Figures S3A-S3C), indicating that FRM-4 and FRM-1 cooperate in maintaining EPJ architecture, with FRM-4 making the larger individual contribution.

To investigate the subcellular localization of FRM-4, we inserted a conditional mStayGold tagging cassette into the endogenous *frm-4* locus and used PVD-specific FLP recombinase to achieve cell-specific fluorescent labeling.^87,88^ Endogenous mStayGold::FRM-4 exhibited a striking patch-like distribution in the PVD soma (Figures 2A and S3D), closely matching the pattern of EPJs. Consistently, mStayGold::FRM-4 showed extensive colocalization with the junctional marker mScarlet::MAPPER (Figure 2C), indicating that FRM-4 is highly enriched at neuronal EPJs.

**Figure 2.**
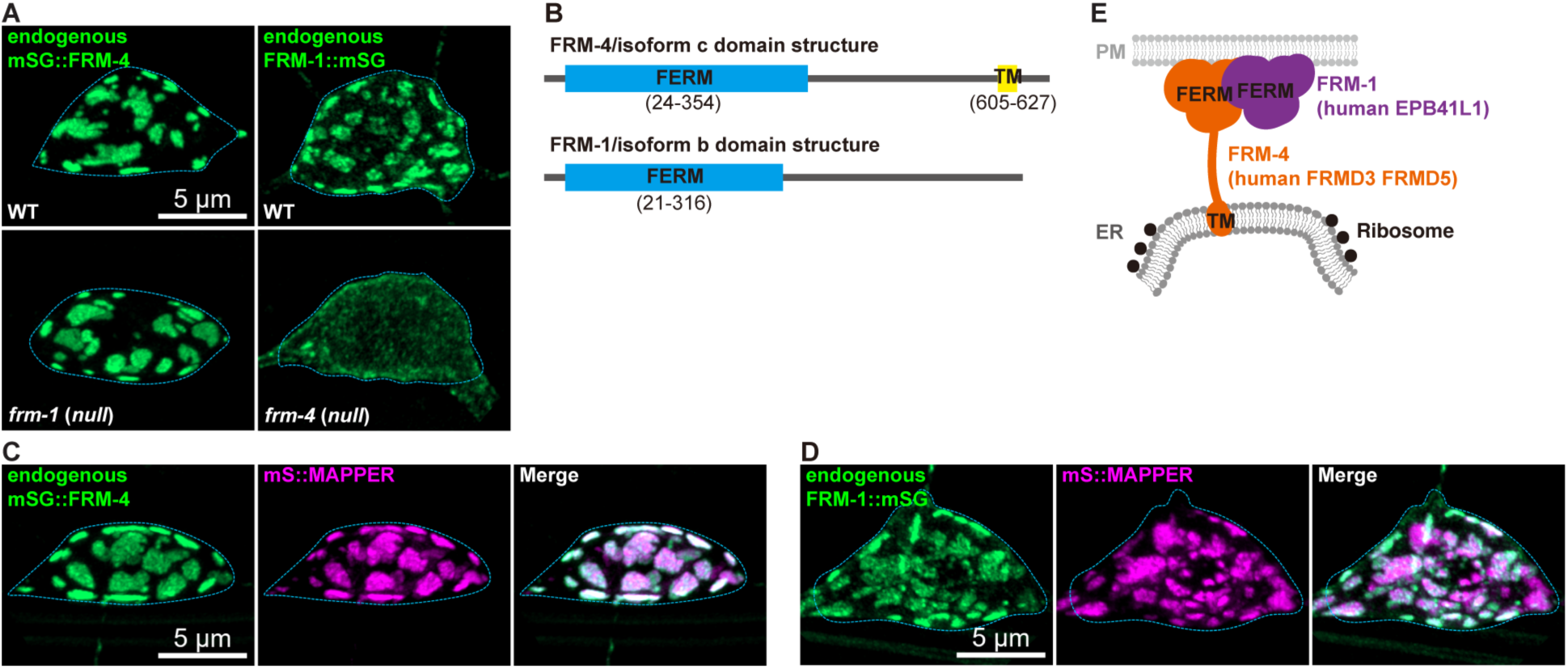
FRM-4 and FRM-1 localize to neuronal EPJs. (A) Representative confocal z-stack projections of endogenously mStayGold-tagged FRM-4 and FRM-1 in PVD somas of WT and indicated mutant animals. (B) Predicted domain organization of FRM-4 and FRM-1, showing their FERM domains and the C-terminal TM segment predicted for FRM-4. (C) Representative confocal z-stack projections showing the spatial overlap of endogenous mStayGold::FRM-4 with mScarlet::MAPPER expressed from a single-copy transgene in the PVD soma. (D) Representative confocal z-stack projections showing the spatial overlap of endogenous FRM-1::mStayGold with single-copy transgenic mScarlet::MAPPER in the PVD soma. (E) Proposed model of FRM-4-dependent FRM-1 enrichment at neuronal EPJs.

FRM-4 contains an N-terminal FERM domain and a predicted C-terminal TM domain but lacks a conventional N-terminal signal peptide (Figure S3F). Based on its domain architecture, we hypothesized that FRM-4 is an ER-resident transmembrane protein localized at EPJs. To investigate the role of its C-terminal TM domain, we generated an endogenous mStayGold-tagged FRM-4 variant lacking this region FRM-4 (ΔTM) and examined its subcellular localization. Deletion of the TM domain markedly reduced the mStayGold::FRM-4 fluorescence signal (Figures S3F and S3G), suggesting that the TM domain contributes to FRM-4 protein stability. Notably, under increased imaging exposure, the residual mStayGold::FRM-4 (ΔTM) signal exhibited a partially plasma membrane distribution rather than the characteristic junctional pattern of wild-type FRM-4 (Figures S3F and S3G). These observations suggest that the TM domain is important for retaining FRM-4 at the ER membrane, whereas its cytosolic region may independently associate with the plasma membrane (Figure S3H).

In PVD neurons, FRM-4 was predominantly localized in the soma and largely absent from neurites, revealed by PVD specific endogenous labeling (Figure S3I). Examination of endogenous FRM-4 expression in other tissues showed a much broader tissue distribution, including neurons, muscle, intestine, hypodermis, germline, oocytes, spermatheca, and embryos (Figure S4A). In non-neuronal tissues, FRM-4 formed punctate structures with varying densities and spatial distributions, whereas neuronal somas displayed the distinctive patch-like pattern (Figure S4A). These observations suggest that FRM-4 is broadly expressed but adopts a particularly organized junctional distribution in neurons.

We next examined the localization of FRM-1. Endogenously tagged FRM-1::mStayGold also formed discrete patches in the PVD soma (Figures 2A and S3E) that colocalized with mScarlet::MAPPER (Figure 2D), indicating that FRM-1, like FRM-4, is enriched at EPJs. However, unlike FRM-4, FRM-1 lacks a predicted TM domain and is therefore expected to be recruited by junctional bridge molecules. To determine whether FRM-4 contributes to FRM-1 recruitment, we examined endogenous FRM-1::mStayGold in *frm-4* (*null*) mutants. Strikingly, loss of FRM-4 abolished the junctional enrichment of FRM-1, resulting in a diffuse distribution near the plasma membrane (Figure 2A). In contrast, deletion of *frm-1* did not appreciably alter the junctional pattern of endogenous mStayGold::FRM-4 (Figure 2A), revealing a hierarchical relationship between the two proteins (Figure 2E).

To determine whether FRM-4 and FRM-1 physically associate, we purified full-length FRM-1 and a FRM-4 (ΔTM) construct lacking its C-terminal TM domain. We first analyzed their oligomeric states using size-exclusion chromatography coupled with multi-angle light scattering (SEC-MALS). Increasing the concentration of either FRM-1 or FRM-4 (ΔTM) caused a modest shift toward smaller elution volumes, suggesting concentration-dependent self-association (Figures S5A and S5B). When the two proteins were mixed, an additional peak appeared at a smaller elution volume than either protein alone (Figure S5C), consistent with the formation of a higher-order FRM-4 (ΔTM) -FRM-1 complex. Pull-down assays further supported a substoichiometric association between FRM-4 (ΔTM) and FRM-1 (Figure S5D). Together, these biochemical observations suggest that FRM-4 and FRM-1 can weakly associate directly. At EPJs, FRM-4 and FRM-1 are likely confined to a crowded 2-dimensional space where weak interactions together through avidity effect can potentially account for FRM-4’s recruitment of FRM-1. Collectively, these findings identify FRM-4 and FRM-1 as cooperative regulators of neuronal EPJ morphology.

### FRM-4 and FRM-1 recruit SOCE machinery to neuronal EPJs

To examine the relationship between FRM proteins and SOCE machinery, we simultaneously visualized endogenously labeled ORAI-1 and FRM-4. Multiple ORAI-1 puncta were positioned within individual FRM-4-enriched patches. STIM-1 showed near complete colocalization with FRM-4 (Figures 3A and 3B). Thus, FRM-4 defines larger junctional territories containing SOCE proteins.

**Figure 3.**
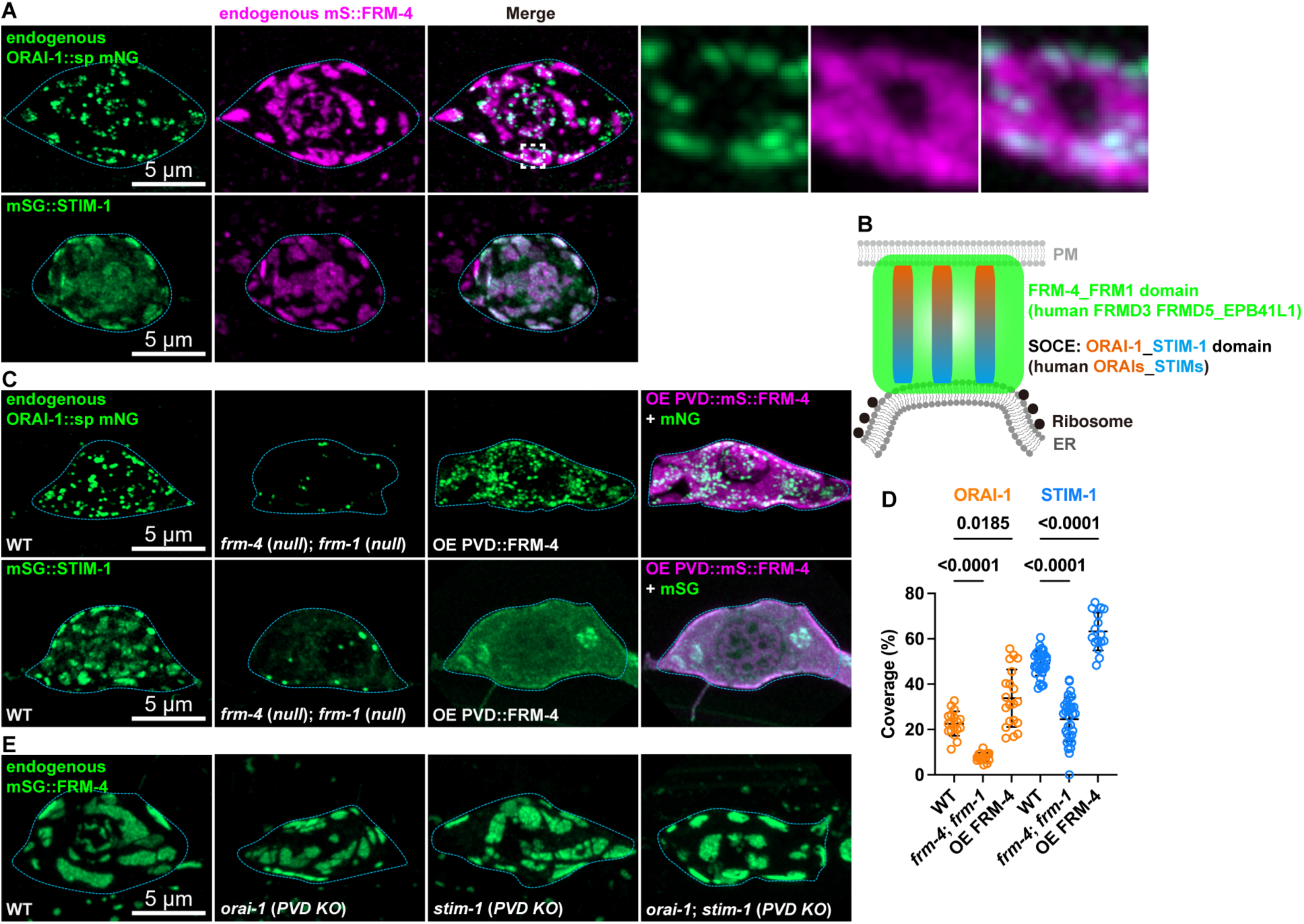
FRM proteins recruit SOCE machinery within junctional territories. (A) Representative confocal z-stack projections showing the spatial relationships of endogenously split-mNeonGreen-labeled ORAI-1 or single copy mStayGold::STIM-1 transgene with endogenously mScarlet-tagged FRM-4 in the PVD soma. (B) Model depicting SOCE-associated puncta within FRM-4/FRM-1-enriched junctional territories. (C) Representative confocal z-stack projections of endogenous split-mNeonGreen-labeled ORAI-1 and single copy mStayGold::STIM-1 transgene in PVD somas of the indicated genotypes. (D) Quantification of ORAI-1 and STIM-1 coverage in the PVD soma of WT, *frm-4*; *frm-1* double-mutant, and FRM-4-overexpressing animals (ORAI-1: n = 18, 17, and 20; STIM-1: n = 33, 33, and 16, respectively). Each dot represents one PVD neuron from an independent animal. Data are presented as mean ± SD. Statistical significance was determined using Welch’s one-way ANOVA followed by Dunnett’s T3 multiple comparisons test. The WT ORAI-1 and STIM-1datasets are the same as those presented in Figure 1G and are reproduced here for comparison. (E) Representative confocal z-stack projections of endogenous mStayGold::FRM-4 in the PVD somas of the indicated genotypes.

In *frm-4*; *frm-1* double mutants, both ORAI-1 and STIM-1 signal was markedly reduced (Figures 3C and 3D), indicating that FRM proteins are required to recruit the SOCE machinery to EPJs. Conversely, overexpression of FRM-4 expanded its distribution across the entire somatic plasma membrane and caused both ORAI-1 and STIM-1 to adopt similarly expanded patterns (Figures 3C and 3D). These findings indicate that FRM proteins are necessary and sufficient to recruit SOCE to EPJs. Moreover, the patch-like pattern and abundance of FRM-4 remained unchanged in *orai-1* and *stim-1* single mutants and in *orai-1*; *stim-1* double mutants, placing FRM proteins upstream of ORAI-1 and STIM-1 in the spatial organization of neuronal EPJs (Figure 3E).

### FRM-4 and FRM-1 pattern CICR machinery at neuronal EPJs

We next investigated how FRM proteins organize CICR machinery, comprising the voltage-gated Ca²⁺ channel EGL-19, the junctional protein JPH-1, and the ryanodine receptor UNC-68. Endogenously labeled CICR components formed puncta along the plasma membrane. Strikingly, these puncta precisely localized to the edges of FRM-4-enriched patches with one puncta per patch (Figures 4A and 4B). This adjacent but mutually exclusive arrangement contrasts with the localization of SOCE components which are localized within FRM-4 territories. This is consistent with the notion that CICR and SOCE proteins occupy distinct but neighboring subdomains of EPJs (Figure 1K). In *frm-4*; *frm-1* double mutants, the numbers of EGL-19, JPH-1, and UNC-68 puncta were markedly reduced (Figures 4C and 4D), indicating that FRM proteins are required for the normal organization of CICR machinery. Notably, FRM-4 overexpression also reduced CICR puncta (Figures 4C and 4D). This result is consistent with the spatial exclusion of CICR components from FRM-4-enriched domains and suggests that FRM mediated-EPJ formation patterns CICR proteins but FRM proteins can also locally excludes the CICR complex.

**Figure 4.**
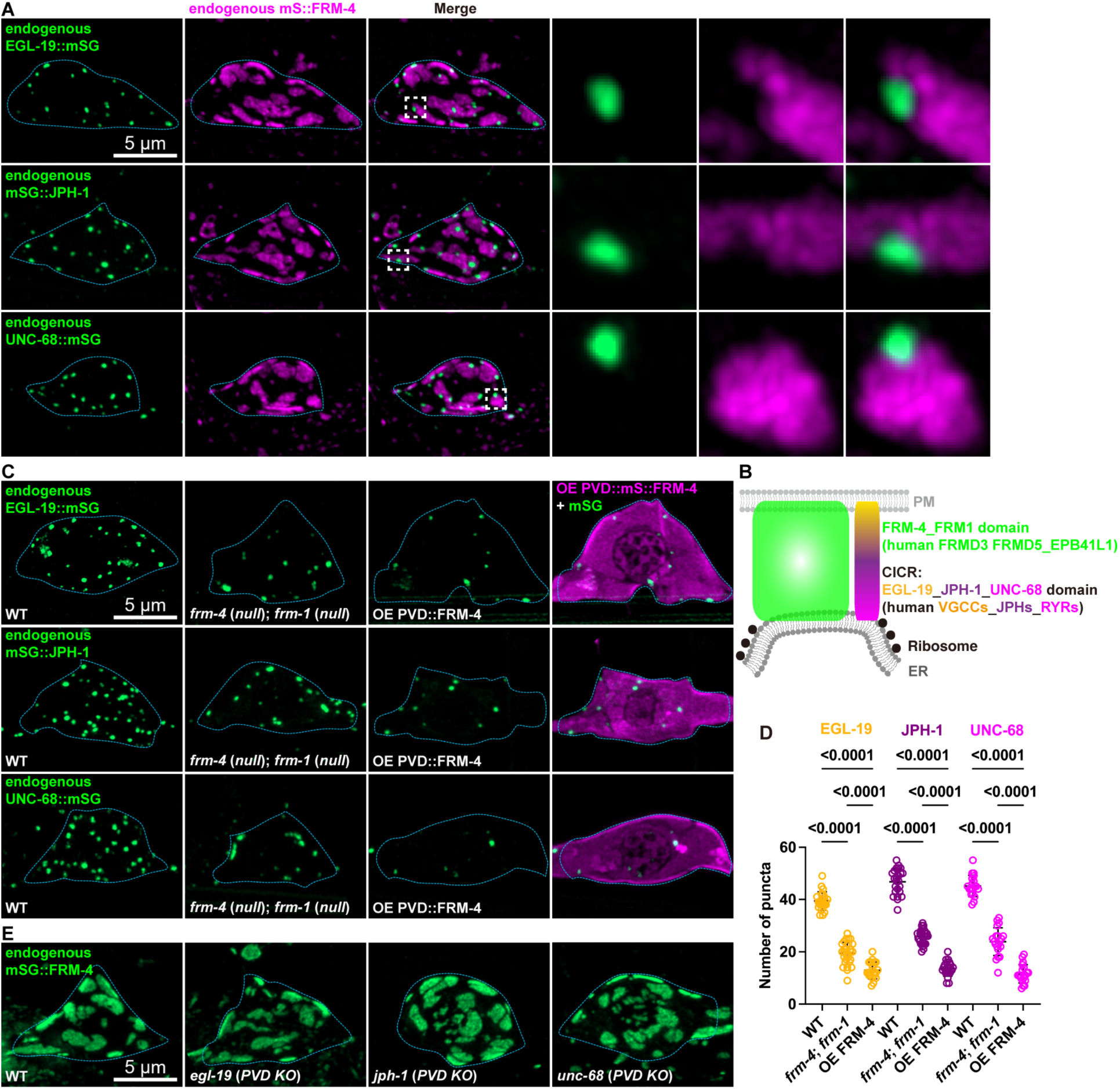
FRM proteins pattern CICR-associated puncta at junctional boundaries. (A) Representative confocal z-stack projections showing the spatial relationships of endogenously mStayGold-tagged EGL-19, JPH-1, or UNC-68 with endogenously mScarlet-tagged FRM-4 in the PVD soma. (B) Model depicting CICR-associated puncta adjacent to FRM-enriched junctional territories. (C) Representative confocal z-stack projections of endogenous mStayGold-tagged EGL-19, JPH-1, and UNC-68 in the PVD somas of the indicated genotypes. (D) Quantification of endogenous EGL-19, JPH-1, and UNC-68 puncta in the PVD neuronal somas of WT, *frm-4* (*null*); *frm-1* (*null*) double-mutant, and single copy PVD FRM-4-overexpressing animals. Each dot represents one PVD neuron from an independent animal. EGL-19: n = 23, 28, and 21; JPH-1: n = 24, 30, and 23; UNC-68: n = 19, 20, and 20, respectively, for WT, double-mutant, and FRM-4 OE animals. Data are presented as mean ± SD. Statistical significance was assessed separately for each protein using one-way ANOVA with multiple-comparisons correction. The WT EGL-19 and UNC-68 datasets are the same as those presented in Figure 1J and are reproduced here for comparison. (E) Representative confocal z-stack projections of endogenous mStayGold::FRM-4 in PVD somas of the indicated genotypes.

Conversely, FRM-4 maintained its patch-like distribution in *egl-19*, *jph-1*, or *unc-68* mutants (Figure 4E), indicating that its localization does not require the CICR machinery. Together, these results place FRM proteins upstream of both SOCE and CICR organization, while revealing distinct spatial relationships with the two signaling modules.

We next asked whether FRM proteins are sufficient to form membrane junctions. To test this, we fused the cytosolic regions of FRM-4 or FRM-1 to a mitochondrial targeting sequence derived from TOMM-20 (Figure S6A). Whereas mitochondria normally occupied the central region of the PVD soma, expression of either fusion protein caused a pronounced redistribution of mitochondria toward the cell cortex (Figure S6B). These findings suggest that the cytosolic regions of FRM-4 and FRM-1 are individually sufficient to promote membrane contact formation.

### FRM-4 and FRM-1 are required for the ultrastructural organization of neuronal EPJs

Next, we asked whether FRM proteins also regulate the membrane architecture of EPJs using cryo-electron tomography (cryo-ET) (Figure S7A). After high-pressure freezing *C. elegans* strains expressing mScarlet in PVD, we used cryo-correlative light and electron microscopy (cryo-CLEM) to locate PVDs within the frozen animals. We then proceeded with lift-outcryo-focused ion beam (cryo-FIB) milling to generate approximately 200-nm-thickness lamellae through the middle section of the PVD soma, followed by cryo-ET data collection and analysis. Consistent with ssTEM observations, tomographic slices and three-dimensional reconstructions revealed closely opposed ER and PM regions (Figures 5A-5C). Mapping local ER-PM distances onto the junction ER surface revealed that separions of about 20-30 nm across most of the EPJs, with a local region of greater separation approaching 50 m. (Figure 5D).

**Figure 5.**
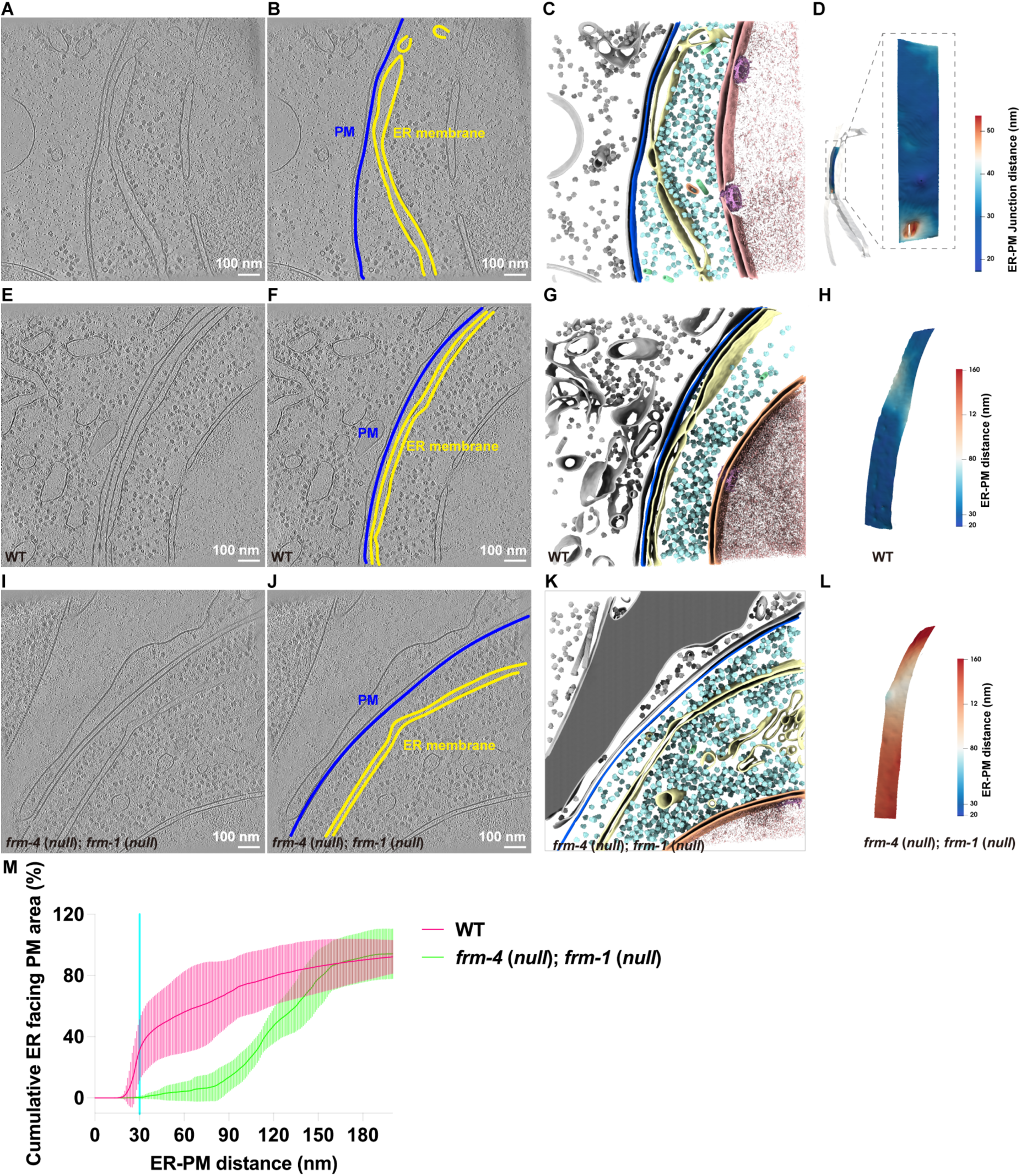
FRM-4 and FRM-1 are required for ER-PM membrane apposition. (A and B) Representative cryo-ET sections through the WT PVD soma showing the PM and cortical ER. (C) Corresponding 3D segmentation of the PM and ER in (A) and (B). (D) 3D rendering of the junctional interface, with an enlarged surface view color-coded by local ER-PM separation. (E and F) Additional representative cryo-ET sections showing an EPJ in the WT PVD soma. (G) Corresponding 3D segmentation of the PM and ER in (E) and (F). (H) 3D rendering of the EPJ color-coded by local ER-PM separation. (I and J) Representative cryo-ET sections through the PVD soma of *frm-4*; *frm-1* double mutants showing the ER-PM interface. (K) Corresponding 3D segmentation of the PM and ER in (I) and (J). (L) 3D rendering of the mutant ER-PM interface color-coded by local membrane separation. (M) Cumulative percentage of the PM-facing ER surface area as a function of the shortest distance to the PM in WT and *frm-4*; *frm-1* PVD neurons. Curves show mean ± SD across eight tomograms per genotype. WT or *frm-4*; *frm-1* n = 8 tomograms from 5 PVD neurons, each PVD neuron from an independent animal.

Compared with wild-type animals, *frm-4*; *frm-1* double mutants, the overall abundance and architecture of ER, PM and other organelles are normal. However, the double mutants exhibited a substantially increased separation between the ER and PM, with numerous ribosomes present in the space between the membranes (Figures 5E-5L). To quantify this defect, we measured the shortest distance from the PM-facing ER surface to the PM and calculated the cumulative fraction of ER surface area within each distance threshold (Figure 5M). In wild-type animals, around 31.3% of the PM-facing ER surface was within 30 nm of the PM, whereas this fraction dropped to only around 0.28% in the double mutants. Thus, FRM proteins are required not only for organizing junctional signaling proteins but also for maintaining close ER-PM membrane apposition.

### FRM proteins and EPJ components undergo phase separation *in vivo* and *in vitro*

Given the weak homophilic and heterophilic interaction between FRM-4 and FRM-1, we wondered how they can form patch-like EPJs and recruit Ca^2+^ signaling components. We first examined the dynamics of junctional proteins in living PVD neurons. Time-lapse imaging revealed that the endogenous FRM-4 patches remained largely stable in position and morphology, although occasionally there are striking events of a cytoskeletal structure (most likely microtubule) transiently poking into a FRM-4 patch (Figure 6A). In these events, a tubule-like clearing forms and then disappears within seconds leaving no scars (Arrows in Figure 6A). This is similar to a photobleaching experiment and suggest that FRM-4 exhibits liquid condensate-like behavior in the patches.

**Figure 6.**
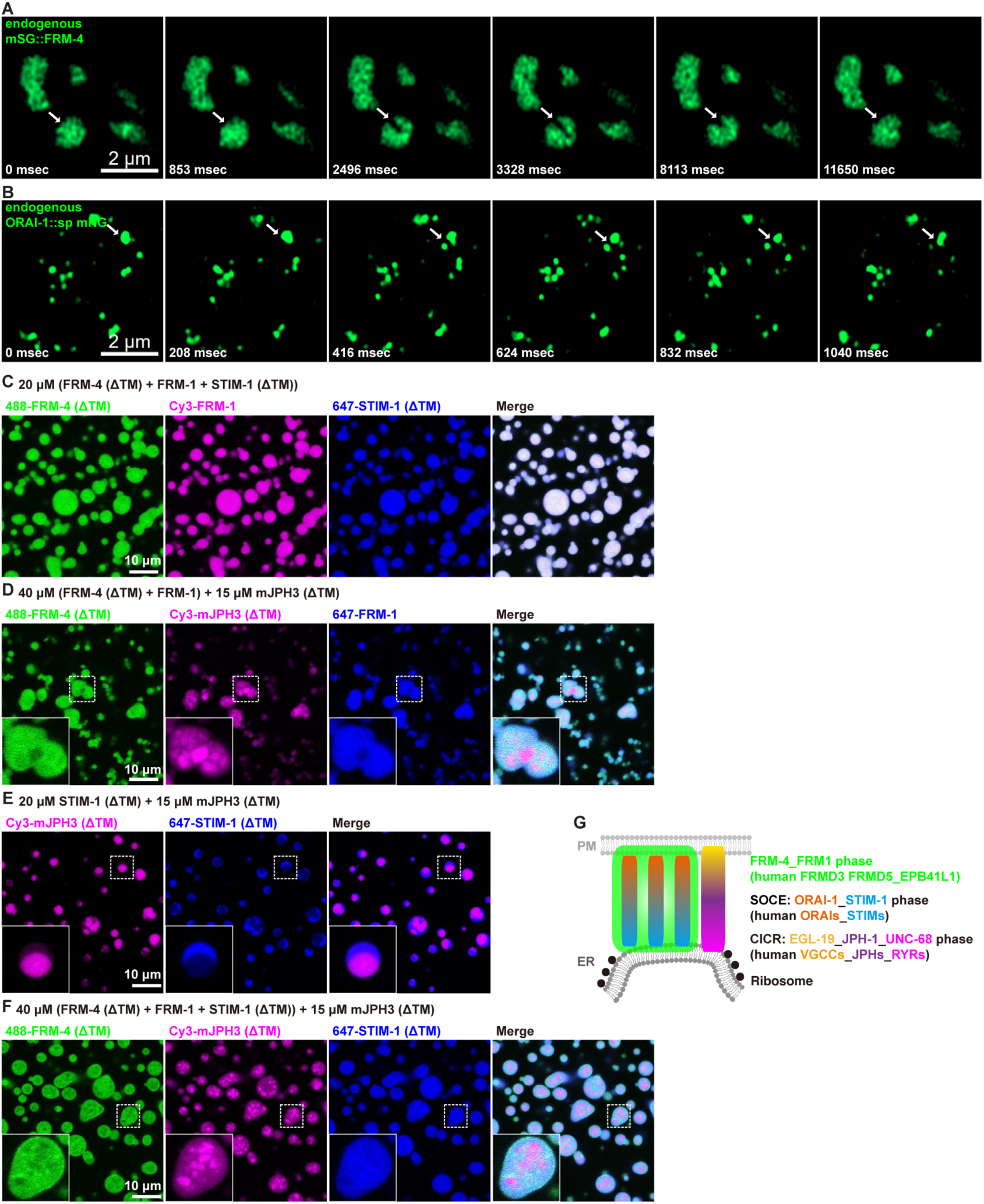
FRM proteins SOCE- and CICR-associated proteins undergo phase separation. (A) Representative single-plane time-lapse confocal images of endogenous mStayGold::FRM-4 in the PVD soma. FRM-4-enriched structures are largely stable, with transient clearing events. (B) Representative single-plane time-lapse confocal images of endogenously split-mNeonGreen-labeled ORAI-1 in the PVD soma, illustrating dynamic puncta. (C) Representative confocal images of condensates formed by purified FRM-4 (ΔTM), FRM-1, and STIM-1 (ΔTM) at the indicated concentrations. Fluorescently labeled protein constituted 1% of each preparation. Unless otherwise indicated, imaging buffer contained 20 mM HEPES (pH 7.5), 100 mM NaCl, 1 mM TCEP, and 3% (w/v) PEG 8000. (D-F) Representative confocal images showing the spatial segregation of mJPH3 (ΔTM) from condensates formed by FRM-1 and FRM-4 (ΔTM) (D), STIM-1 (ΔTM) (E), or FRM-1, FRM-4 (ΔTM), and STIM-1 (ΔTM) (F), at the indicated protein concentrations. Insets show threefold magnifications of representative condensates. (G) Model illustrating selective partitioning of SOCE- and CICR-associated proteins into adjacent domains at neuronal EPJs; phase separation is proposed as an organizing mechanism.

Different from FRM-4, endogenously labeled ORAI-1 puncta were highly dynamic, displaying frequent fusion and fission events within seconds (Figure 6B; Video 1). Such behavior is consistent with the notion that ORAI-1 form dynamic molecular assemblies which undergo LLPS at EPJs. Dual color imaging further showed that ORAI-1 puncta moved within FRM-4-enriched patches while largely remaining confined to those territories (Video 2). These results suggest that the ORAI-1 condensates are immisive within the larger FRM-4 condensates *in vivo*.

In contrast, CICR-associated puncta were comparatively stable over time, although occasional fusion- or fission-like events were observed (Video 3). During junctional remodeling, CICR puncta remained associated with the edges of FRM-4 domains and are excluded from the FRM-4 patch rather than mixing with their interiors (Video 4). These findings reveal distinct dynamic behaviors of the two signaling modules and suggest that FRM-4 tethered junctional regions support ORAI-1 condensates but exclude CICR proteins, leading to the spatial separation of two Ca^2+^ signaling complexes.

We next tested whether FRM proteins could form condensates in a reconstituted system. Purified FRM-4 (ΔTM) and FRM-1 each formed concentration-dependent, liquid-like assemblies *in vitro* (Figures S8A-S8C), supporting their intrinsic capacity for LLPS. We next examined how FRM proteins influence the partitioning of SOCE components within condensates. STIM-1 (ΔTM) formed condensates on its own (Figure S9A). When mixed with FRM-1 and FRM-4 (ΔTM), STIM-1 (ΔTM) was incorporated into a shared condensate phase (Figure 6C). Interestingly, removal of FRM-1 resulted in partial demixing of STIM-1 (ΔTM) and FRM-4 (ΔTM), which remained within the same droplets but exhibited distinct phases (Figure S9C). Consistent with this observation, STIM-1 (ΔTM) fully mixed with FRM-1 condensates in the absence of FRM-4 (Figure S9B). These results suggest that FRM-1 promotes the mixing of STIM-1 and FRM-4 within condensates.These observations indicate that the composition of FRM assemblies determines their capacity to incorporate STIM-1 and is consistent with the genetic results that both FRM proteins are required for EJP formation.

We next tested the mammalian CICR-associated protein mouse mJPH3, a homolog of *C. elegans* JPH-1. mJPH3 (ΔTM) formed condensates *in vitro* (Figure S9D) but remained spatially separated from either FRM-1 or FRM-4 (ΔTM) condensates in pairwise mixtures (Figures S9E and S9F). mJPH3 (ΔTM) also segregated from condensates containing both FRM proteins (Figure 6D). Consistent with the separation of SOCE and CICR components *in vivo*, mJPH3 (ΔTM) and STIM-1 (ΔTM) formed distinct condensate phases when mixed together (Figure 6E).

Finally, when FRM-4 (ΔTM), FRM-1, STIM-1 (ΔTM), and mJPH3 (ΔTM) were combined, STIM-1 partitioned with the FRM-containing phase, whereas mJPH3 (ΔTM) remained enriched in a separate phase (Figures 6F and 6G). Together, these reconstitution experiments yielded remarkably similar results compared to *in vivo* experiments and demonstrate that FRM-containing condensates can differentially partition components associated with SOCE and CICR. This protein sequence based, selective mixing and demixing provides a potential biophysical mechanism for the neighboring but mutually exclusive subdomains observed at neuronal EPJs *in vivo*.

### FRM-4 and FRM-1 are required for EPJ function in vivo

To assess the physiological importance of FRM-dependent EPJ organization, we measured the mechanical stimulus-induced cytosolic Ca^2+^ response in PVD soma.^89^ We trapped worms in microfluidic chambers and repeatedly applied defined harsh touch stimuli while recording GCAMP signals in the soma (Figures 7A and 7B). This paradigm allowed us to measure the magnitude of individual evoked responses as well as the progressive changes in Ca²⁺ signaling across repeated stimulation. In wild-type animals, successive stimuli elicited Ca²⁺ transients that progressively increased in amplitude, accompanied by an accumulating residual Ca²⁺ signal (Figures 7A and 7B).

**Figure 7.**
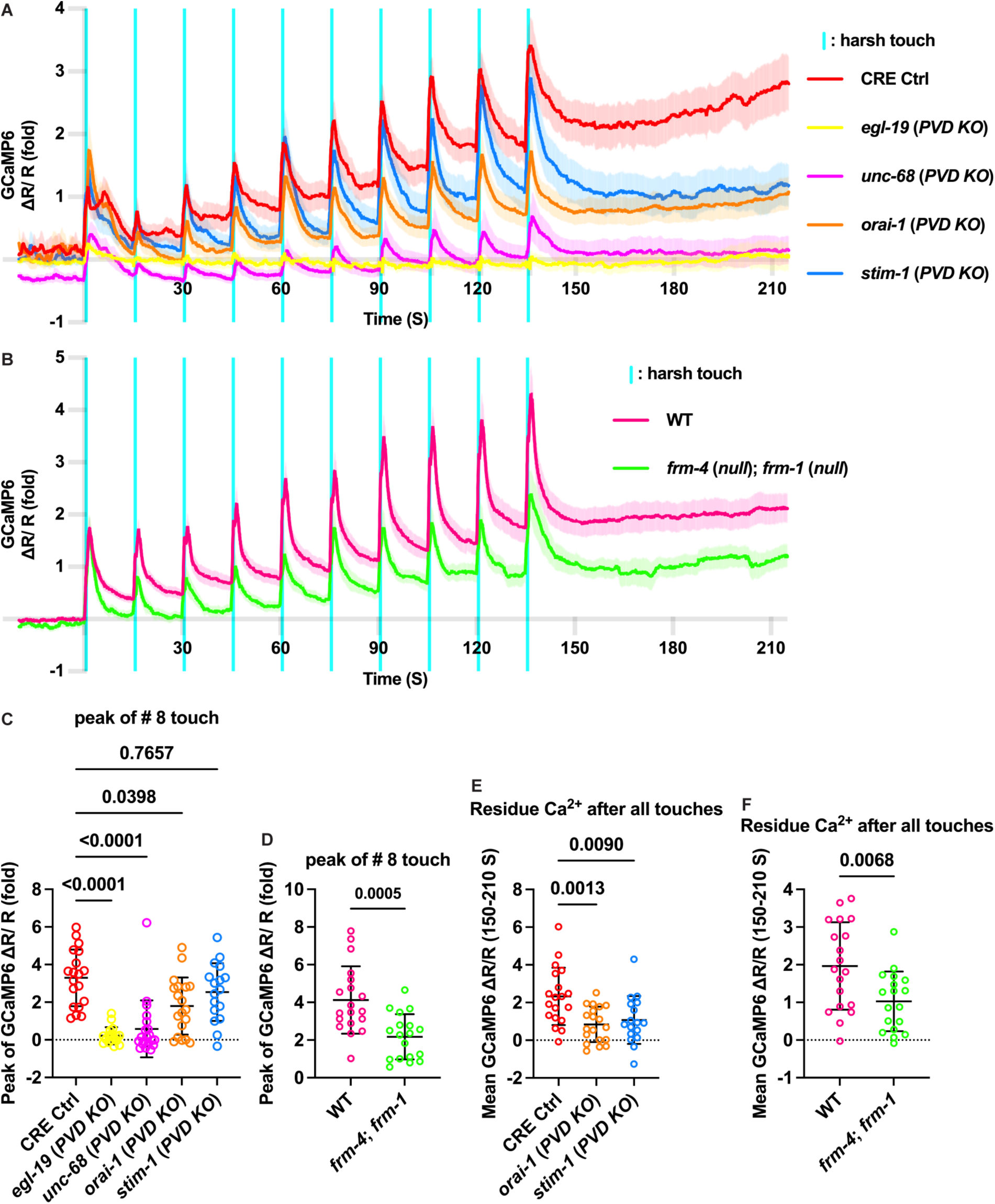
EPJ proteins support different aspects of stimulus-evoked Ca²⁺ signals. (A and B) Average GCaMP6 fluorescence responses (ΔR/R) in PVD somata following repeated harsh-touch stimulation (40 psi), cyan vertical lines indicate the timing of harsh touch stimuli. Shaded regions or error bars represent SEM. (A) Cre control (n = 19) and PVD-specific knockouts of *egl-19* (n = 18), *unc-68* (n = 20), *orai-1* (n = 19), and *stim-1* (n = 17). (B) WT (n = 19) and *frm-4* (*null*); *frm-1* (*null*) double mutants (n = 18). (C and D) Quantification of peak GCaMP6 responses to the eighth harsh touch stimulus for the indicated genotypes in (A) and (B), respectively. Each dot represents one PVD neuron from an individual animal. (E and F) Quantification of sustained Ca²⁺ responses, measured as the mean GCaMP6 ΔR/R during the 150-210 s interval following the final harsh touch stimulus, for the indicated genotypes in (E) Cre control, *orai-1* (*PVD KO*), and *stim-1* (*PVD KO*) and (F) WT and *frm-4* (*null*); *frm-1* (*null*). Each dot represents one PVD neuron from an individual animal. Data are presented as mean ± SD in (C-F). Statistical significance was determined using Brown-Forsythe and Welch’s one-way ANOVA followed by Dunnett’s T3 multiple comparisons test (C), two-tailed unpaired Welch’s t test (D and F), or ordinary one-way ANOVA followed by Dunnett’s multiple comparisons test (E). Exact p values or p-value thresholds are indicated.

To distinguish the contributions of CICR and SOCE to these responses, we examined mutants affecting each pathway. Loss of the CICR components EGL-19 or UNC-68 cause a dramatic reduction of the peak Ca²⁺ responses to individual touches, suggesting that EGL-19 is primarily responsible for the initial Ca^2+^ entry which is then amplified by Ca²⁺ release from ER through UNC-68 (Figures 7A, 7C, S10A). On the contrary, *orai-1* and *stim-1* mutants exhibited normal peak responses, suggesting that the slow Ca²⁺ entry from ORAI-1 does not significantly contribute to individual Ca²⁺ spike (Figures 7A, 7C, S10A). At the end of each stimulus, Ca²⁺ level does not go back to the baseline, indicative of insufficient Ca²⁺ clearing. *orai-1* and *stim-*1 mutants show a trend toward reduced residual Ca²⁺ after individual spike. However, statistical significance was observed only at 6-10^th^ individual spike in *orai-1* mutants (Figure S11A). Nevertheless, both *orai-1* and *stim-*1 mutants showed significantly reduced residue cytosolic Ca²⁺ after all 10 stimuli (Figure 7E), suggesting that the accumulative Ca²⁺ increase is partially dependent on SOCE. These results demonstrate that the CICR and SOCE machineries are responsible for distinct aspects of Ca²⁺.

In *frm-4*; *frm-1* double mutants, the individual Ca²⁺ spikes are lower in their amplitude, consistent with the reduced CICR proteins at EPJs (Figures 7B, 7D, S10B).

Furthermore, the *frm-4*; *frm-1* double mutants exhibited reduced residual Ca²⁺ signals (Figures 7B, 7F, S11B). Together, these findings indicate that FRM proteins support both the individual evoked Ca²⁺ responses through recruitment of CICR proteins and the progressive accumulation of residual Ca²⁺ during repeated sensory stimulation through SOCE pathway (Figures S12).

### FRM-dependent organization of neuronal EPJs is evolutionarily conserved

Finally, we asked whether the spatial organization observed in *C. elegans* neurons is conserved in human neurons. When overexpressed in iPSC induced human neurons, the human FRM-4-related proteins FRMD3 and FRMD5 formed prominent patch like structures at the cell surface (Figures S13A and S13B). By contrast, the FRM-1-related protein EPB41L1 did not exhibit the same patch-like distribution (Figure S13C).

The SOCE-associated protein STIM2^26^ displayed both patch-like structures and a broader ER-associated pattern (Figure S13D), whereas the CICR-associated protein JPH3 also formed discrete junctional structures (Figure S13E). Dual-color imaging revealed substantial overlap between FRMD3 or FRMD5 and STIM2 (Figures S13F and S13H). In contrast, FRMD3- and FRMD5-enriched regions were predominantly adjacent to JPH3-positive structures (Figures S13G and S13I). Thus, SOCE- and CICR-associated proteins exhibit distinct spatial relationships with FRM-family proteins in human neurons, reminiscent of their organization in *C. elegans*. Together, these observations suggest that aspects of FRM-associated spatial organization at neuronal EPJs are evolutionarily conserved.

## Discussion

Our study identifies two nested levels of organization of EPJs in the neuronal cell body. At the cellular level, EPJs form discrete patches across the soma rather than a continuous interface. Within these patches, proteins associated with SOCE and CICR occupy neighboring but distinct subdomains. FRM-4, the central organizer of the patch-like EPJs, is largely restricted to the neuronal soma and the immediate proximal dendrite, suggesting that the patch-like structure is a somatic specialization. Consistent with this notion, patch-like EPJs are seen in other *C. elegans* neuron somas by serial sectioning electron microscopy and FRM-4-related proteins display a patch-like distribution in human iPSC-derived neurons (Figures S1 and S13). FRM-4 is expressed broadly in many tissues and adopts diverse subcellular localization patterns (Figure S4), suggesting that the organization of EPJs is tuned in a cell type-specific manner.

FRM-4 and FRM-1 are central to this organization. Loss of FRM function reduces the number of junctional patches, diminishes the junctional enrichment of both signaling modules and results in greater distance between ER membrane and PM. FRM proteins therefore contribute both to the protein composition and to the physical apposition of the junction. The two proteins appear to have distinct roles. FRM-4 carries a C-terminal transmembrane segment that retains it in the ER membrane, whereas its cytosolic FERM-containing region can associate with the plasma membrane on its own, as suggested by the FRM-4 (ΔTM) variant. FRM-1 lacks a transmembrane segment and depends on FRM-4 for its junctional enrichment. FERM domains classically link membrane proteins, phosphoinositides, and the cortical cytoskeleton,^86,90^ so identifying the PM-side partners that anchor FRM-4 across the gap is an important next step.

Purified FRM-4 and FRM-1 interact only weakly, yet FRM-1 recruitment *in vivo* strictly requires FRM-4. We propose that confining both proteins to the narrow cleft between two membranes, at high local concentration and reduced dimensionality, converts weak multivalent interactions into stable, avidity-driven assemblies.^91–93^ Consistent with a scaffolding role, FRM-4 localization is independent of ORAI-1, STIM-1, EGL-19, JPH-1, and UNC-68, its overexpression expands both its own territory and that of SOCE components, and the cytosolic regions of FRM-4 and FRM-1 are each sufficient to pull mitochondria toward the cortex. It is also possible that a dense FRM assembly sterically excludes ribosomes from the apposed ER surface, accounting for the ribosome-free zone that defines EPJs in our ssTEM data, although this remains to be tested.

Weak, multivalent protein interactions can drive concentration-dependent LLPS,^57,94^ and several of our observations suggest that FRM-organized EPJs exhibit condensate-like properties. Most studies of biomolecular condensates rely on overexpression in cultured cells or on purified proteins.^94^ Here we examined endogenously tagged proteins in the neurons of living animals, which allowed us to follow the relative behavior of the different phases in their native context. FRM-4 territories are relatively stable in shape and position, yet are cleared reversibly and without a trace when a cytoskeletal filament transiently passes through them, as expected for a fluid-like scaffold. ORAI-1 puncta fuse and divide within FRM-4 territories while remaining confined to them, and CICR puncta remain at the territory edges and are excluded from its interior even as the junction remodels. Thus, a large FRM-4 phase contains a smaller, dynamic, immiscible ORAI-1 phase and abuts a third CICR phase that does not mix with either. This is the type of multiphase organization described for nucleolar sub-compartments and other multi-component condensates,^95,96^ and here we observe the phase relationships directly in intact neurons at endogenous expression levels rather than inferring them from *in vitro* mixtures.

The reconstitution experiments provide a complementary test of sufficiency. FRM-4 (ΔTM) and FRM-1 each form condensate *in vitro.* STIM-1 (ΔTM) mixes with FRM-1 but demixes from FRM-4 (ΔTM) alone, and is brought into a shared phase only when FRM-1 is added. In contrast, mJPH3 (ΔTM) remains in a separate phase from FRM condensates and from STIM-1 under all conditions tested. These results suggest that composition tunes miscibility: FRM-4 nucleates the territory, FRM-1 acts as a bridge that makes it compatible with SOCE components, and CICR components are excluded by their own intrinsic interaction preferences. These data are aligned with the genetic necessity of both FRM-1 and FRM-4, as well as the localization patterns of CICR and SOCE.

Condensate assisted-segregation of molecules according to their function has been reported in epigenetic regulation. Chromatin regulators with opposing activities are sorted into separate condensates. HP1 proteins compact nucleosomes into droplet-like heterochromatin that excludes much of the transcription machinery,^65,66^ whereas transcription factors, Mediator, and RNA polymerase II concentrate at active enhancers in distinct condensates.^97,98^ Nested sub-compartments within the nucleolus similarly arise from sequence-encoded differences in miscibility.^95^ The EPJ shares several features with these systems. Functionally distinct activities, here Ca²⁺ release from the ER and Ca²⁺ reentry into it, are physically separated without any surrounding membrane. The FRM-4/FRM-1 condensate defines the territory, and neighboring territories remain adjacent but do not mix. There are also important differences. EPJ condensates are anchored to two apposed membranes and are effectively two-dimensional, so geometry as well as sequence is likely to shape the pattern, for example the single CICR punctum we observe at the edge of each FRM-4 patch. In addition, the two compartments in chromatin carry out largely opposing programs, whereas SOCE and CICR draw on a shared ER Ca²⁺ store and must remain functionally coupled. Adjacency without mixing may therefore be the key design principle. It places Ca²⁺-release and Ca²⁺-reentry machinery right next to one another, while keeping their regulation independent. The latter may be important because STIM1 has been reported to inhibit Ca_V_1.2 channels,^42,43^ so spatial segregation could prevent cross-inhibition between the two pathways. This idea remains speculative and will require direct testing.

The relationship between CICR and SOCE is also important from a functional perspective. In neurons, Ca²⁺ influx through L-type channels can be amplified by ryanodine receptor-mediated release from the ER, with junctophilins coupling the two channel types.^49,99^ This amplification shapes membrane excitability, for example through Ca²⁺-activated K⁺ channels, and activity-dependent gene expression.^49,51^ SOCE is generally slower and smaller in amplitude in neurons, and has been implicated in refilling ER stores and sustaining prolonged Ca²⁺ signals, in spine stability, and in synaptic plasticity.^40,100–102^ Our results fit this division of labor. Loss of EGL-19 or UNC-68 strongly reduces the peak response to each touch, consistent with Ca_V_1-mediated entry amplified by ER release. In contrast, loss of ORAI-1 or STIM-1 leaves peak responses intact but reduces the accumulation of residual Ca²⁺ across repeated stimuli. We speculate that SOCE helps maintain the ER store and thereby sustains the progressive amplification of Ca²⁺ responses seen in wild-type neurons, a form of short-term sensitization that is lost when SOCE or the FRM scaffold is compromised. An attractive implication of the adjacent architecture is that Ca²⁺ entering through ORAI-1 would be pumped into the ER close to nearby ryanodine receptors, allowing local refilling of the release sites.^38,103^ Direct measurements of ER Ca²⁺ dynamics in these neurons will be needed to test this model.

Our findings in human neurons raise the possibility that this organization is conserved. The human FRM-4-related proteins FRMD3 and FRMD5 form patches at the cell surface that overlap with STIM2 and sit adjacent to JPH3, in a manner reminiscent of the *C. elegans* soma, although these experiments rely on overexpression and the segregation is less pronounced than in worms. Patchy ER-PM contacts have also been described in mammalian neurons, for example those organized by Kv2.1 channels and VAP proteins,^104,105^ suggesting that discrete, protein-defined junctions are a general feature of neuronal somata. The implications of this organization may extend to disease. Variants in FERM-domain-containing genes, including FRMD7, FRMD4A, and FRMD5, have been associated with neurological and neurodevelopmental disorders.^106–108^ Similarly, JPH3 repeat expansions cause Huntington disease-like 2,^50^ while dysregulation of ryanodine receptors, STIM-dependent store-operated Ca²⁺ entry, and L-type Ca²⁺ channel signaling has been implicated in neurodegenerative disease.^40^ Whether disrupted junctional organization contributes to these conditions, and whether it affects neuronal resilience to stress or aging, is an open question that this system is well suited to address.

## Resource Availability

### Lead contact

Requests for further information and reagents should be directed to the lead contact, Kang Shen, at, Huichao Deng, at.

## Materials availability

Plasmids generated in this study will be available in the Addgene repository.

## Acknowledgments

We thank all members of the Shen laboratory for insightful discussions and constructive feedback. We are grateful to Hang Lu, Nathan McDonald, and members of their laboratories for their support and assistance to Huichao in conducting the harsh-touch stimulation assays. We thank Liqun Luo for providing access to an upright confocal microscope for functional testing experiments. We thank the Lavis lab and Open Chemistry team (Janelia) for the gift of JF552-cpSNAP dye. We also thank Jeremy Magescas from the Feldman laboratory for providing the *C. elegans* codon-optimized mStayGold plasmid.

K.S. is an Investigator of the Howard Hughes Medical Institute. This work was supported in part by the Knight Initiative for Brain Resilience at Stanford University, Shenzhen Medical Research Fund (E250200111) to X.W., the Fundamental and Interdisciplinary Breakthrough Plan of the Ministry of Education of China (JYB2025XDXM505 to M.Z. and X.W.).

## Author Contributions

Conceptualization, H.D. and K.S.; Methodology, H.D., J.C., R.F., X.T., A.C., W.H., G.Q., X.Li., C.W., R.D., J.Z., S.K., M.Z., H.L., M.L., X.W., and K.S.; Investigation, H.D. (*C. elegans* related experiments), J.C., M.Z., and X.W. (*in vitro* phase separation assays), R.F. (serial-section transmission electron microscopy), X.T. and M.L. (cryo-electron tomography), H.D., A.C., and H.L. (harsh-touch assays), H.D., W.H., and X.Li. (human neuron experiments), H.D., G.Q. (image analysis), H.D., C.W., R.D., and S.K. (functional assays), and J.Z. (structural prediction), X.L. and C.T. provided tool reagents on *C. elegans*, J.F. provided *C. elegans* mStayGold plasmid; Resources, H.D., and K.S.; Writing - Original Draft, H.D. and K.S.; Writing - Review & Editing, H.D. and K.S., with input from all authors; Supervision, K.S.; Funding Acquisition, K.S.

## Declaration of Interests

The authors declare no competing interests.

## STAR★Methods

### C. elegans strains

*C. elegans* strains were grown on OP50 *E. coli*-seeded nematode growth media (NGM) plates at 20°C, following standard protocols.^109^ N2 Bristol is the wild-type reference strain, and all the mutants were isolated from N2. All animals were day1 adults unless otherwise indicated.

All the endogenous knock in worms were generated by CRISPR/Cas9 editing.^79^ Transgenic strains were generated using standard microinjection techniques.

All the *C. elegans* strains used in this study are listed in table S1.

### Molecular biology

Plasmids, crRNAs, primers sequence used to generate transgenic or knock in *C. elegans* strains or human neurons in this study are listed in table S1. Plasmids were generated using In-fusion cloning kit. crRNAs used for CRISPR/Cas9 knock in were ordered from IDT.

### Serial-section transmission electron microscopy

Worms were prepared for conventional EM by high pressure freezing/freeze-substitution. Worms in M9 containing 20% BSA and *E. coli* were frozen in 100 μm well specimen carriers (Type A) opposite a hexadecane coated flat carrier (Type B) using a Leica EM ICE high-pressure freezer (Leica Microsystems, Deerfield, IL). Freeze-substitution in 1% OsO4, 0.1% uranyl acetate, 1% methanol in acetone, containing 3% water^110,111^ was carried out with a Leica AFS2 unit. Following substitution, samples were rinsed in acetone, infiltrated and then polymerized in Eponate 12 resin (Ted Pella, Inc, Redding, CA). Serial 40 nm sections were cut with a Leica UC7 ultramicrotome using a Diatome diamond knife, picked up on Pioloform coated slot grids and stained with uranyl acetate and Sato’s lead.^112^ Montage imaging of sections was performed with an FEI Tecnai T12 TEM at 120 kV using a Gatan 4k x 4k Rio camera at a nominal magnification of 4400x (pixel size 1.54nm) using SerialEM. TrakEM2 in Fiji was used to align serial sections montages into a 3D volume.^113–115^ Modeling of 3D volumes was performed with IMOD.^116^

### Microscopy for C. elegans and human neurons

*C. elegans* and human neurons fluorescence imaging was performed on a Zeiss LSM 980 Airyscan 2 system with a 63× Plan-Apochromat 1.4NA objective and Hoechst 33342, GFP, mNeonGreen, mScarlet lasers, Alexa Fluor 647 laser. The Airyscan 4Y multiplex mode was used to capture super resolution images. Animals were imaged live on 5% agarose pads in 10 mM levamisole for imaging.

### Image analysis and quantification

Airyscan images and analysis were processed with Zeiss ZEN software. Statistical calculations and graphing were done in Prism 11 (GraphPad).

For every 3D fluorescence PVD ORAI-1, STIM-1 image, we estimate a proxy surface by fitting the cell-body signals. More specifically, we use Li thresholding to identify the cell body and radial edges to locate the boundary of the cells. The fitted shape is represented as a triangle mesh. The bright regions on cell membranes are then segmented with different methods. For ORAI-1 signals, we use signal-to-noise thresholding. For STIM-1, a method that combines multi-Otsu thresholding with local contrast recovery is used to retain puncta. We use signed-distance interpolation to estimate the intersections between the triangles and the bright mask. For the STIM-1 signals, we further use surface-normal projection to map bright signals near the fitted surface onto the mesh. Finally, we sum the bright areas on the cell membrane and divide them by the reference surface area to obtain the coverage of bright areas.

### Protein expression and purification

FRM-1 full-length, FRM-4 (ΔTM), STIM-1 (ΔTM), and Mouse JPH-3 (ΔTM) (aa 1-724, mJPH3) genes were cloned into a modified pET32a vector, with a Trx-6xHis tag and a protease 3C cutting site at the N-terminal of the multi-clone sites. Sequences were confirmed by sequencing. Proteins were expressed in Escherichia coli BL21-CodonPlus(DE3)-RIL cells (Agilent). Cells were grown in LB at 37°C with 220 rpm shaking to an OD600 of ∼ 0.8, followed by 0.5 mM IPTG induction for protein expression. Continue to culture the cells overnight for protein expression. Collect the cell pellet by centrifuging at 4,200 rpm for 10 min at 4°C. Resuspend the pellet with binding buffer composed of 50 mM Tris pH 8.0, 1000 mM NaCl, 5 mM Imidazole, supplemented with 1 mM PMSF. Homogenize the suspension with a French press high-pressure homogenizer, followed by centrifugation at 18,000 rpm for 30 min at 4°C. The supernatant was collected and drain through a gravity Ni2+-NTA Sepharose (GE Healthcare) column, washed with binding buffer three times, and eluted with 50 mM Tris pH 8.0, 500 mM NaCl, 300 mM imidazole. Then the proteins were subject to size-exclusion chromatography with HiLoad 26/600 Superdex 200pg column, pre-equilibrated with 50 mM Tris pH 8.0, 100 mM NaCl, 1 mM EDTA, 1 mM DTT. Fractions with proper proteins were pooled and digested by 3C protease overnight at 4°C. An anion or cation exchange column was used to remove the Trx-6xHis tag and other impurities for FRM-1 and FRM-4, respectively. Then the proteins were loaded to another round of HiLoad 26/600 Superdex 200pg column equilibrated with HBS (HEPES buffer saline: 20 mM HEPES pH 7.5, 100 mM NaCl, 1 mM TCEP). Proteins were concentrated with an Amicon ultra centrifugal filter (MWCO = 10 kDa, Millipore) to ∼ 10 mg/ml and aliquoted into small fractions. Proteins were then flash frozen in liquid nitrogen and stored at - 80°C.

### Size exclusion chromatography coupled with multiangle light scattering (SEC-MALS)

The SEC-MALS system is composed of a multi-angle light scattering (MALS) detector (miniDawn, Wyatt), a differential refractive index (dRI) detector (Optilab, Wyatt), and a Liquid chromatography (LC) system (AKTA pure, GE Healthcare). For each experiment, 200 μl sample was loaded via injection loop (100 μl) into a Superdex 200 10/300 GL column (GE Healthcare) equilibrated with HBS. Data were analyzed by ASTRA6 (Wyatt).

### Fluorescent labelling for in vitro proteins

Fluorescent labeling dyes iFluor 488/Cy3/Cy5 NHS esters (AAT Bioquest) were dissolved in DMSO to a stock concentration of 10 mM. Proteins were concentrated to 5 ∼ 10 mg/ml mass concentration in HBS before labeling. Mix the protein and dye in a 1:1 molar ratio. Keep the mixture in dark with gentle shaking at room temperature for 1 h. The reaction was stopped by adding 200 mM Tris. Labeled proteins were centrifuged to remove any precipitation, then loaded onto a HiTrap desalting column (GE Healthcare) equilibrated with HBS. Fluorescence labeling efficiency was determined by NanoDrop OneC (Thermo scientific). For imaging assays, sparse labeling was used by diluting the labeled proteins with unlabelled ones.

### Microscopy for in vitro proteins

Protein aliquots were centrifuged at 21,300 g for 10 min at 4°C to remove potential aggregates before performing experiments. Proteins were diluted with HBS buffer to desired concentrations. PEG 8000 (if needed) at the indicated concentrations was added after the proteins were well mixed in the buffer. Samples were injected into a homemade chamber with 0.15 mm thickness. For multiple-component phase separation assays containing STIM-1, a passivated chamber was used to maintain a relatively round shape of the condensate. The chamber was sealed immediately with vaseline after sample loading. After 15 minutes of incubation, the samples were imaged with a Zeiss LSM 980 confocal microscope with a 63x oil lens. All imaging experiments were completed within 1.5 h after sample mixing.

### High-pressure freezing (HPF) sample preparation of C. elegans for cryo-ET

Synchronized *C. elegans* were washed off NGM plates using M9 buffer, and the resulting suspension was transferred into 1.5 mL tubes. The suspension was centrifuged at ∼1500 rpm (400×g) for 1-2 min, and the pellet containing worms was incubated in 10 mM levamisole (prepared in M9 buffer) for 5 min for anesthetization. The following HPF procedure was optimized Waffle method:^117^ Anesthetized nematodes were loaded onto 50-mesh copper grids coated with a formvar film. The grids were placed into 6-mm-diameter high-pressure freezing carriers (Leica Microsystems). A drop of 60% Ficoll in M9 buffer was applied to the grid to fully immerse the worms, followed by sealing the sample with a sapphire disc. The assembled sample was rapidly placed into the central well of the middle plate; spacers were added on top of the sapphire disc to fill the height of the well. The assembly was then transferred to a Leica EM ICE high-pressure freezer (Leica Microsystems) for rapid freezing. Frozen samples were stored in liquid nitrogen until further processing.

### Correlative light and electron microscopy and lift-out cryo-FIB milling

To precisely locate the PVD neurons within vitrified *C. elegans* samples embedded in the ice block on grids prior to lift-out cryo-focused ion beam (cryo-FIB), cryo-CLEM was performed. The frozen samples on grids were loaded into a plasma cryo-SEM/FIB system (Thermo Scientific Helios™ 5 Hydra CX DualBeam) equipped with an iFLM module and an EasyLift nanomanipulator. The PVD of *C. elegans* was imaged using the iFLM correlative imaging system at excitation wavelengths of 470 nm (JPH-1::mStayGold) and 565 nm (PVD::myr-mScarlet), respectively. The fluorescent images were subsequently registered with the corresponding SEM and ion beam images using MAPS software (Thermo Scientific). Based on the identified location of the PVD, lift-out cryo-FIB milling was performed to generate thin lamellae.

Lift-out cryo-FIB milling was performed with our previously optimized procedure.^118^ A sample block with PVD neuron was milled at 4 nA with Xenon source, and then lifted out with EasyLift nanomanipulator. Next, the block was attached on a pre-milled receiver grid with six-sided attachment method, generating a rough lamella with a height of about 5 um containing the central PVD. Subsequently, to perform fine milling, the front and back surfaces of the lamella were milled to expose clean surfaces, a layer of metal-organic platinum was sputtered for extra protection, and milling was conducted at a stage tilt of 7°: rough milling was performed sequentially at 1 nA to 2.5 µm, 0.3 nA to 1.5 µm, and 0.1 nA to 0.8 µm, with final polishing at 30 pA to generate a lamella approximately 200 nm thick.

### Cryo-ET data collection and reconstruction

Tomographic tilt series acquisition was performed using a Titan Krios G4 (Thermo Fisher Scientific) electron microscope operated at 300 kV, equipped with a Falcon 4i direct electron detector. Tilt series were collected with SerialEM^119^ and PACE-tomo software.^120^ The tilt series were acquired in movie mode using the EER file format at nominal magnifications of 42,000× (pixel size, 2.97 Å) and 15,000× (pixel size, 8.52 Å). Tilting was performed from + 60° to - 60°, starting from a predetermined milling angle, with a tilt increment of 3°, resulting in a total dose of 120 e-/Å2. The target defocus range was set between -4 and -6 μm.

Tilt series were processed using Aretomo software. Then high quality tilt series were further manually aligned with IMOD software^121^ to generate reconstructed tomograms at a binning factor of 4. These tomograms were further denoised with Icecream software to increase image contrast for subsequent visualization and analysis.^122^ Structure visualization and movie generation were conducted with UCSF ChimeraX.^123^

### Quantification of ER-PM proximity from cryo-ET

The ER and PM segmentation was initially performed using Membrain v2^124^ and subsequently refined with Amira (Thermo Fisher Scientific). Surface meshes were generated using Surface Morphometrics.^125^ PM-facing ER was defined as the complete side of the ER membrane facing the PM, including regions outside close contacts. Morphologically apparent WT ER-PM junctions were additionally isolated to characterize junction spacing.

ER-PM distances were calculated using custom Python scripts as the shortest distance from each ER triangle center to the PM polygonal surface. Local measurements were weighted by ER triangle area to calculate cumulative distance distributions and distance percentiles. Junction spacing was summarized by the area-weighted median distance (D50) and the D25-D75 interval. For genotype comparisons, ER-PM proximity was quantified using the fraction of PM-facing ER area within 30 nm of the PM (C30) and the D50 of the complete PM-facing ER surface. Threshold sensitivity was assessed over 20-50 nm. 3D distance maps were visualized in ParaView (2024 kitware, inc.), and figures were prepared using OriginPro 2024 (OriginLab).

### Harsh touch assay

Harsh touch experiments were performed using a microfluidic mechanical stimulation system.^89^ Worms expressing cytosolic GCaMP6 and mCherry were loaded into the imaging channels of the microfluidic devices and loosely held in place with 1 psi of pressure. Stimuli were delivered by applying 40 psi of pressure to PDMS membranes spanning both sides of the area 100-250 µm posterior to the most anterior part of the worm. For all trials, we recorded simultaneous red and green fluorescence of the nearest PVD soma for a 30 s pre-stimulus period followed by 10 bouts of 1 s stimulus every 15 s.

### Calcium imaging for harsh touch assay

Harsh touch calcium imaging was performed using a Leica DMI3000 B inverted microscope with a 40x air objective (NA 0.60) and a dual-band GRP/mCherry filter set (Chroma 59022). Simultaneous two-channel image sequences were captured using a Hamamatsu Flash 4.0 v2 sCMOS camera with 50 ms exposure time using a DV2 beamsplitter (Photometrics) containing a GFP/RFP filter. Excitation light was provided by a Spectra X light engine (Lumencor) triggered by camera exposure.

In each recording frame, the PVD soma was identified in the red channel as the brightest point after difference of Gaussians filtering. A 50 by 50 pixel region of interest around this point was cropped and Otsu thresholded to provide a foreground mask. A background mask was also obtained as the difference of two successive dilations of the foreground mask with a 5-pixel-radius disk. Red-channel signal was measured as the mean foreground pixel intensity minus the mean background pixel intensity. Green-channel signal was measured the same way from the matching ROI in the green channel using the masks obtained from the red channel. Calcium activity is reported as the ratio of green-channel signal to red-channel signal, normalized to the value before the first stimulus.

### Human iPSC and iPSC-derived neuronal cultures

Human induced pluripotent stem cell (iPSC) research was performed with approval from the Stanford Stem Cell Research Oversight (SCRO) board. KOLF2.1J PB-TO-oNGN2 iPSCs^126^ were cultured at 37°C, 5% CO2 in StemFlex media (Thermo Fisher Scientific) on Matrigel-coated wells, and passaged 1:20 every 3-4 days with ReLeSR (STEMCELL Technologies).

Cortical neurons were generated using doxycycline-inducible expression of NGN2.^126^ In brief, iPSCs were plated on Matrigel-coated wells in Induction Media (DMEM/F-12 HEPES with 1x GlutaMAX, 1x NEAA, 1x N2, and 2 µg/mL doxycycline) on Day 0. On Day 3, cells were replated on poly-D-lysine laminin double-coated wells in Maturation Media (50% DMEM/F-12 HEPES : 50% BrainPhys, with 1x N21-MAX, 10 ng/mL BDNF, GDNF, and NT-3, 1 µM uridine and FUDR, 1 µg/mL mouse laminin, and 2 µg/mL doxycycline). Half-volume media changes with Maturation Media were performed on Days 4, 7, 10, and 14. From day 7 onwards, mitotic inhibitors were removed. From Day 10 onwards, Maturation Media contained 100% BrainPhys (STEMCELL technologies). Neurons were transfected with plasmids on Day 10 using Lipofectamine Stem Transfection Reagent (Thermo Fisher).

### Immunocytochemistry

iPSC-derived neurons were fixed with 2% paraformaldehyde (PFA, Electron Microscopy Services) for 5 min, followed by 4% PFA in PBS for 10 min at room temperature. Neurons were permeabilized in 0.1% Triton X-100 in PBS for 5 min, and blocked in 10% goat serum in PBS, stained with MAP2 primary antibody (PhosphoSolutions, 1:2000), then Alexa Fluor 647 secondary antibody (Thermo Fisher), counterstained with Hoechst 33342, and mounted on glass slides in Aqua/Poly-Mount (Polysciences).

## Supplemental video and Excel table titles and legends

### Video S1. Dynamic remodeling of ORAI-1 puncta in the PVD soma, related to Figure 6

Time-lapse imaging of endogenously labeled ORAI-1 reveals highly dynamic puncta exhibiting frequent fusion- and fission-like events in living PVD neurons.

### Video S2. Dynamic ORAI-1 puncta remain confined within FRM-4-enriched domains, related to Figure 6

Simultaneous time-lapse imaging of endogenously labeled ORAI-1 and FRM-4 reveals that ORAI-1 puncta undergo dynamic rearrangements within relatively stable FRM-4-enriched domains while largely remaining confined to these territories.

### Video S3. CICR-associated puncta exhibit limited dynamic remodeling in the PVD soma, related to Figure 6

Time-lapse imaging of endogenously labeled calcium-induced calcium release (CICR) components reveals predominantly stable puncta, with occasional fusion- and fission-like events.

### Video S4. JPH-1 puncta remain associated with the boundaries of FRM-4-enriched domains, related to Figure 6

Simultaneous time-lapse imaging of endogenously labeled JPH-1 and FRM-4 reveals that JPH-1 puncta remain preferentially positioned along the boundaries of FRM-4-enriched domains during junctional remodeling.

### Table S1. C. elegans strains, Plasmids, crRNAs, Primers used in this study

During the preparation of this work, the authors used ChatGPT (OpenAI) and Claude (Anthropic) to improve the clarity, readability, and language of the manuscript and to assist with data analysis. The authors reviewed and verified the outputs, made revisions as needed, and take full responsibility for the content of the published article.

## Notes

### Competing Interest Statement

The authors have declared no competing interest.

