## supplemental figures for "Phase-separated FRM proteins create signaling subdomains in inter-organelle junctions"

### Document S1. Figures S1-S13

#### Contents

Figure S1. Somatic EPJs form discrete patches in *C. elegans* head neurons, related to Figure 1
Figure S2. SOCE and CICR components form distinct somatic junctional patterns, related to Figure 1
Figure S3. FRM-4 and FRM-1 are required for EPJ patterning, related to Figure 2
Figure S4. FRM-4 is expressed in multiple tissues, related to Figure 2
Figure S5. Purified FRM-4 and FRM-1 exhibit both homophilic and heterophilic interaction *in vitro*, related to Figure 2
Figure S6. FRM-4 and FRM-1 promote membrane contact formation, related to Figure 4
Figure S7. Cryo-ET workflow for *C. elegans* sample preparation, related to Figure 5
Figure S8. Purified FRM-4 and FRM-1 undergo LLPS *in vitro*, related to Figure 6
Figure S9. Purified STIM-1 and mJPH3 exhibit selective mixing with FRM condensates *in vitro*, related to Figure 6
Figure S10. Peak $Ca^{2+}$ responses to repeated harsh-touch stimulation, related to Figure 7
Figure S11. Residual $Ca^{2+}$ signals following repeated harsh-touch stimulation, related to Figure 7
Figure S12. Proposed model for FRM-dependent organization of $Ca^{2+}$-signaling subdomains at neuronal EPJs
Figure S13. FRMD3 and FRMD5 exhibit distinct spatial relationships with STIM2 and JPH3 in human neurons

**A**

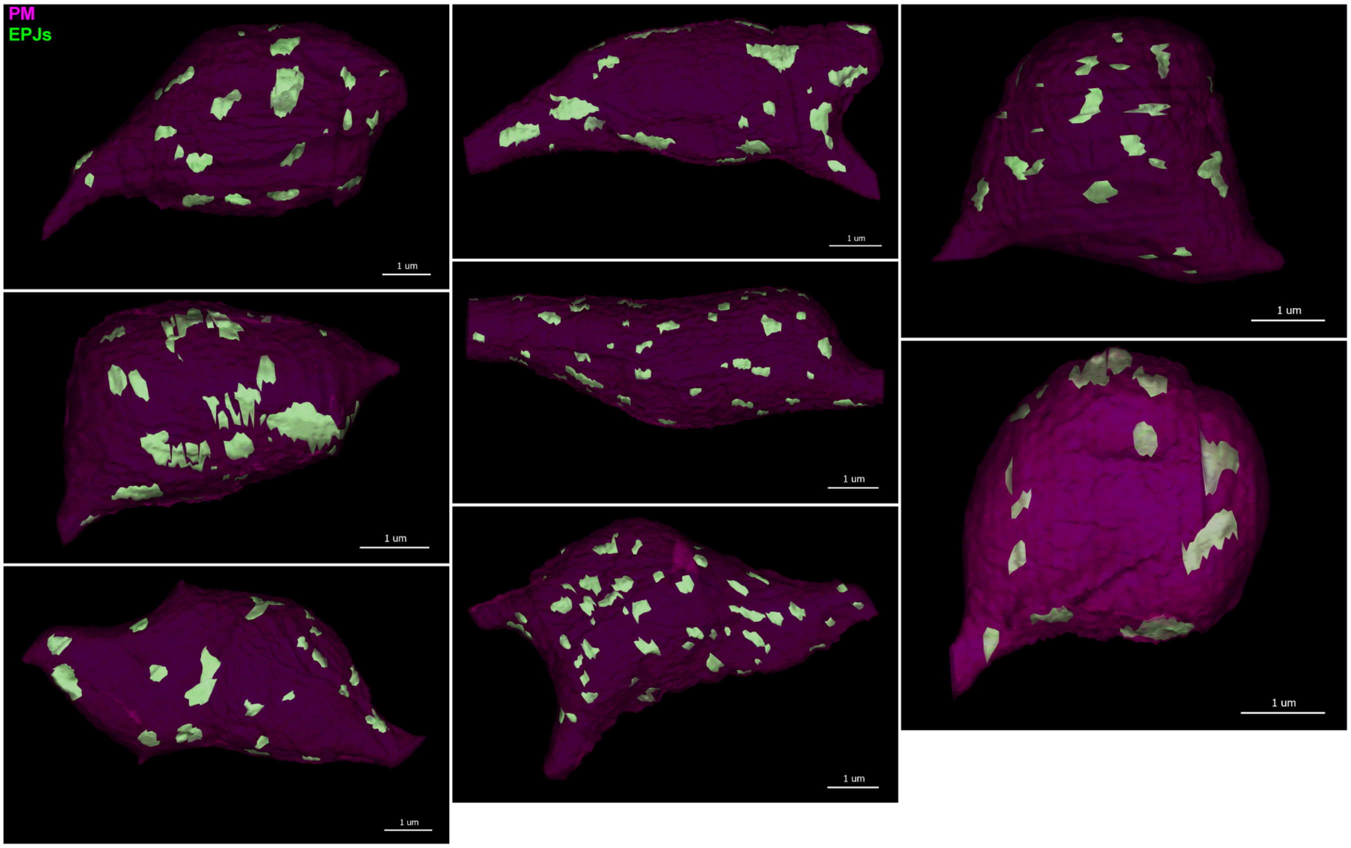


**Figure S1. Somatic EPJs form discrete patches in *C. elegans* head neurons, related to Figure 1**

(A) 3D reconstructions of head neuron somas from ssTEM, showing the spatial distribution of EPJs. EPJs were identified using the same ultrastructural criteria described in Figure 1.

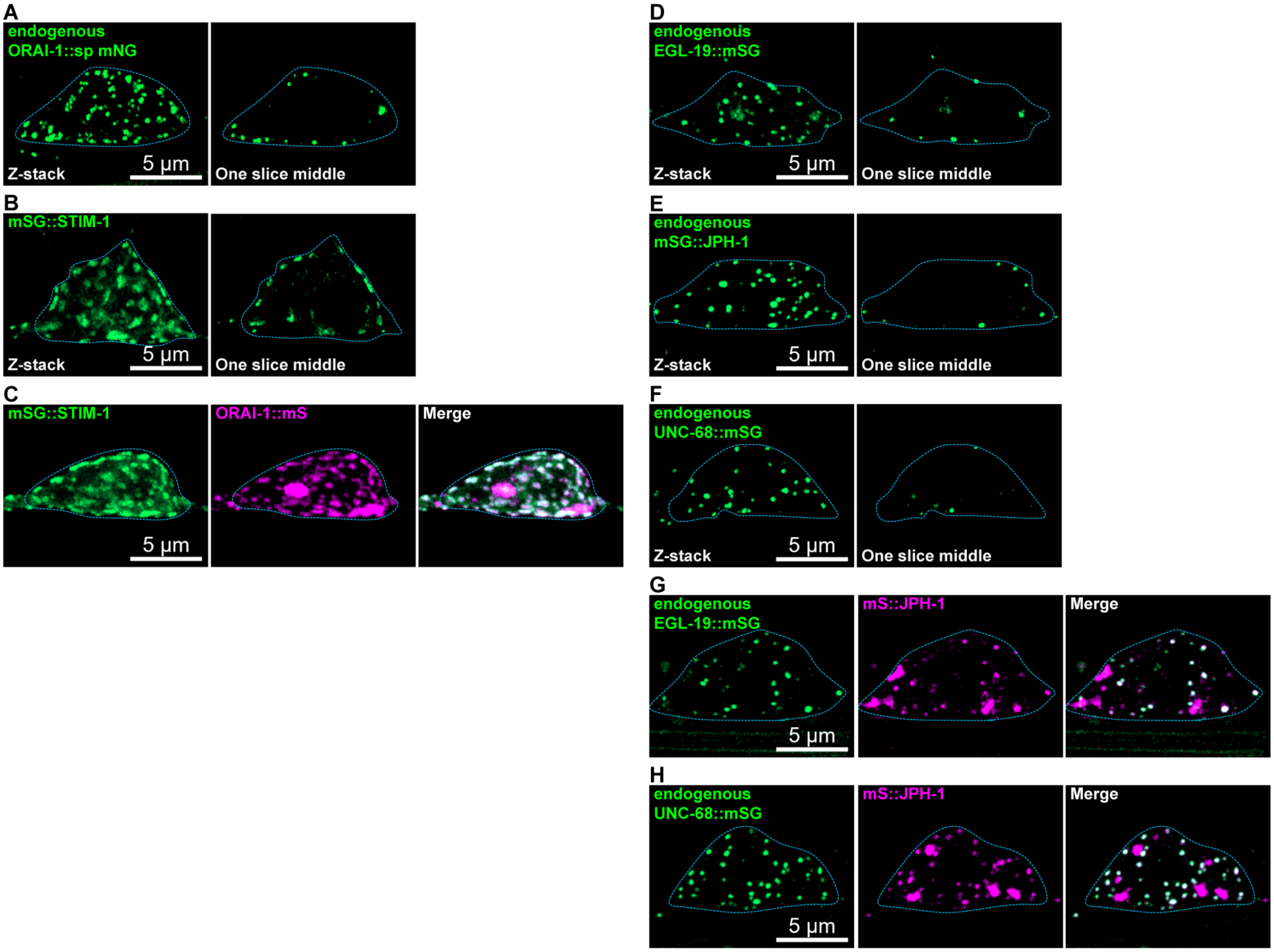


**Figure S2. SOCE and CICR components form distinct somatic junctional patterns, related to Figure 1**

(A) Representative confocal z-stack projections and single optical sections showing endogenously split mNeonGreen-labeled ORAI-1 in the PVD soma.

(B) Representative confocal z-stack projections and single optical sections showing single copy mStayGold::STIM-1 transgene in the PVD soma.

(C) Representative confocal z-stack projections showing the colocalization of mStayGold::STIM-1 and ORAI-1::mScarlet expressed from single-copy transgene in the PVD soma.

(D) Representative confocal z-stack projections and single optical sections showing endogenously mStayGold-tagged EGL-19 in the PVD soma.

(E) Representative confocal z-stack projections and single optical sections showing endogenously mStayGold-tagged JPH-1 in the PVD soma.

(F) Representative confocal z-stack projections and single optical sections showing endogenously mStayGold-tagged UNC-68 in the PVD soma.

(G) Representative confocal z-stack projections showing the colocalization of endogenously mStayGold-tagged EGL-19 and mScarlet::JPH-1 expressed from a single-copy transgene in the PVD soma.

(H) Representative confocal z-stack projections showing the colocalization of endogenously mStayGold-tagged UNC-68 and mScarlet::JPH-1 expressed from a single-copy transgene in the PVD soma.

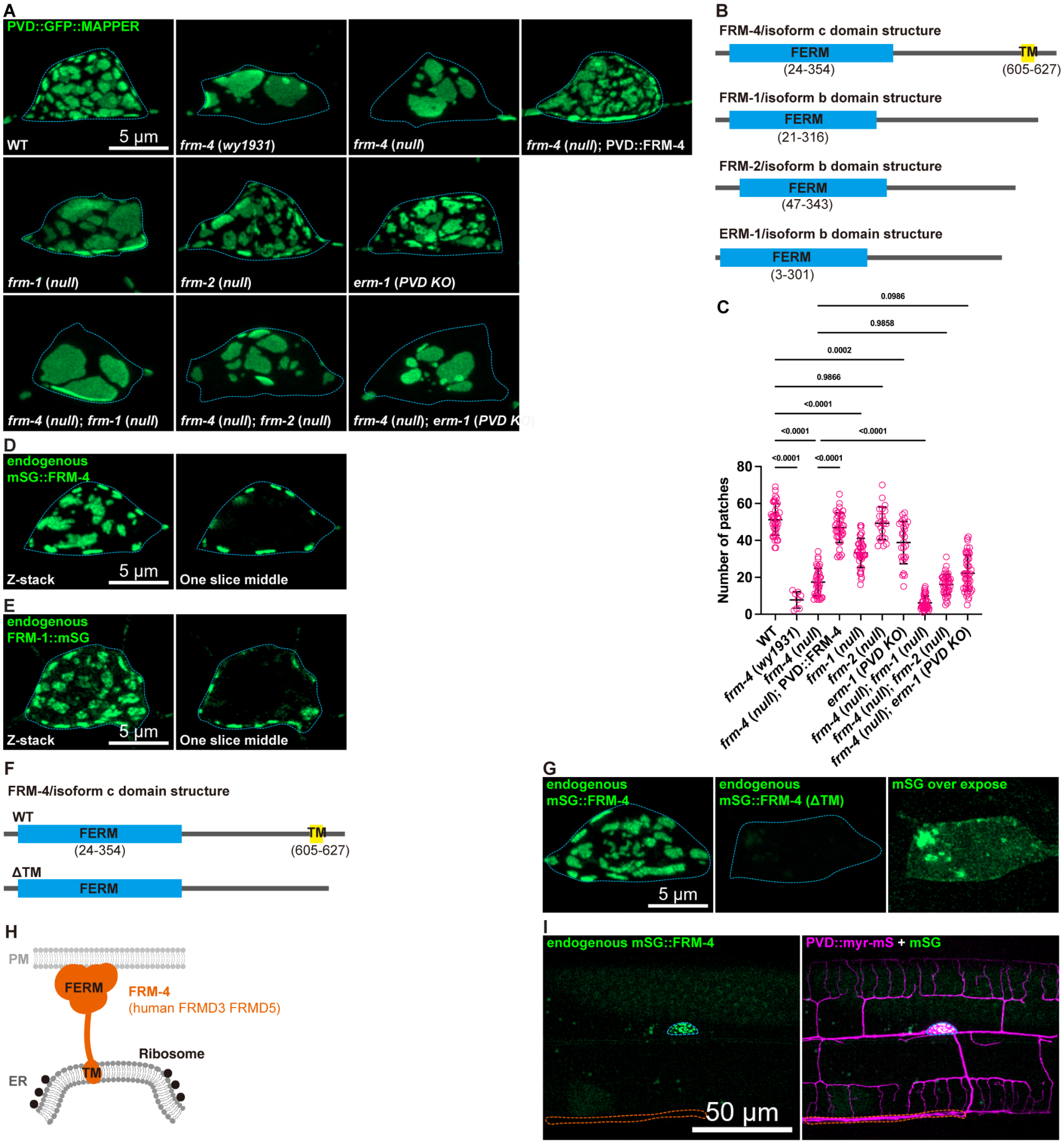


**Figure S3. FRM-4 and FRM-1 are required for EPJ patterning, related to Figure 2**

(A) Representative confocal z-stack projections showing the distribution of GFP::MAPPER in the PVD soma of WT and mutant animals.

(B) Schematic representations of the predicted domain architectures of FRM-4, FRM-1, FRM-2, and ERM-1, highlighting their conserved FERM domains and the predicted C-terminal TM domain of FRM-4.

(C) Quantification of the number of GFP::MAPPER patches per PVD soma in the indicated genotypes. Each dot represents one PVD neuron from an independent animal. Data are presented as mean ± SD. Statistical significance was assessed using Welch's one-way ANOVA followed by Dunnett's T3 multiple-comparisons test. WT, n = 40; *frm-4* (*wy1931*), n = 8; *frm-4* (*null*), n = 38; *frm-4* (*null*); PVD::FRM-4, n = 39; *frm-1* (*null*), n = 39; *frm-2* (*null*), n = 22; *erm-1* (*PVD KO*), n = 26; *frm-4* (*null*); *frm-1* (*null*), n = 41; *frm-4* (*null*); *frm-2* (*null*), n = 41; *frm-4* (*null*); *erm-1* (*PVD KO*), n = 46.

(D and E) Representative confocal z-stack projections and single optical sections showing endogenously mStayGold-tagged FRM-4 (D) and FRM-1 (E) in the PVD soma.

(F) Schematic representations of the predicted domain architectures of full-length FRM-4 and FRM-4 (ΔTM), highlighting the deletion of the C-terminal TM domain.

(G) Representative confocal z-stack images showing the subcellular distribution of endogenously mStayGold-tagged full-length FRM-4 and FRM-4 (ΔTM) in the PVD soma.

(H) Schematic model illustrating the proposed membrane association and organization of FRM-4 at neuronal EPJs.

(I) Representative confocal z-stack projections showing the spatial relationship between endogenously mStayGold-tagged FRM-4 and the plasma membrane marker myr-mScarlet expressed from a single-copy PVD-specific transgene.

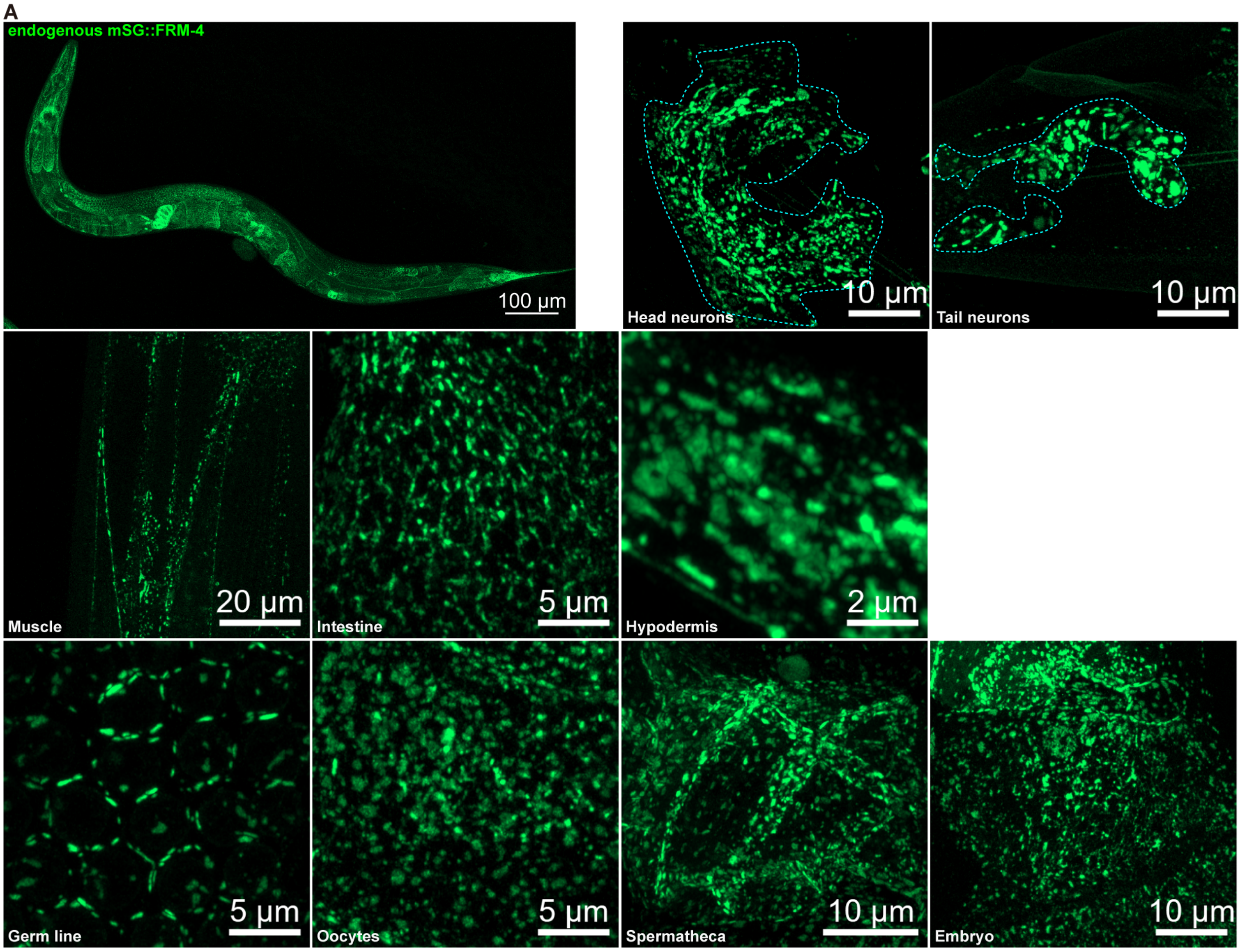


**Figure S4. FRM-4 is expressed in multiple tissues, related to Figure 2**

(A) Representative confocal z-stack projections showing the distribution of endogenously mStayGold-tagged FRM-4 in additional neurons and non-neuronal tissues.

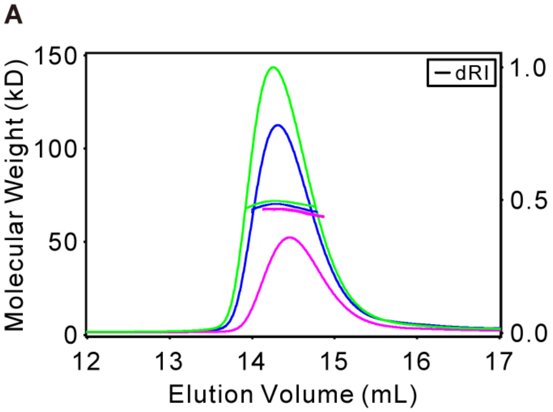


| FRM-1 | Theoretical MW | Measured MW | $V_{elution}$ (ml) |
|---|---|---|---|
| 40 μM | 72.1 kD | 66.4 ± 0.3 kD | 14.46 |
| 80 μM | 72.1 kD | 69.0 ± 0.8 kD | 14.31 |
| 108 μM | 72.1 kD | 70.9 ± 0.5 kD | 14.25 |

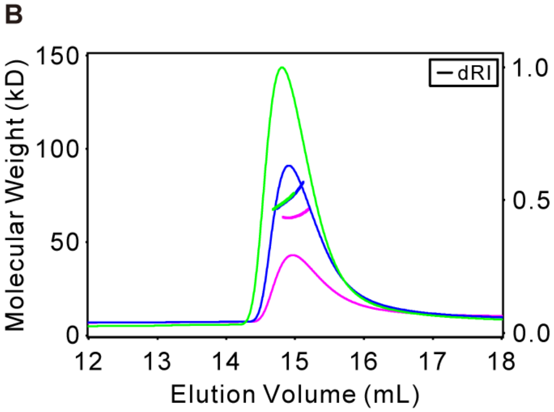


| FRM-4 | Theoretical MW | Measured MW | $V_{elution}$ (ml) |
|---|---|---|---|
| 40 μM | 64.6 kD | 64.5 ± 9.0 kD | 14.97 |
| 80 μM | 64.6 kD | 74.2 ± 19.1 kD | 14.91 |
| 120 μM | 64.6 kD | 71.6. ± 7.1 kD | 14.81 |

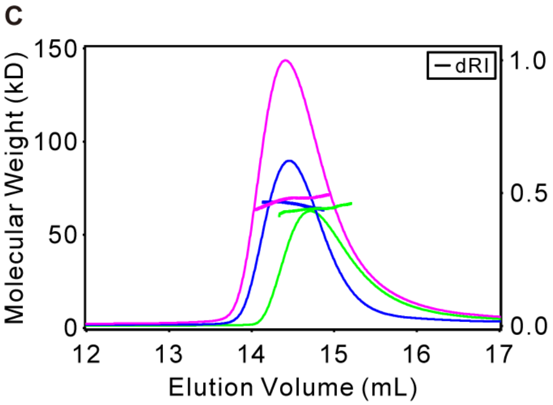


| | Theoretical MW | Measured MW | $V_{elution}$ (ml) |
|---|---|---|---|
| FRM-4 | 64.6 kD | 64.0 ± 3.6 kD | 14.70 |
| FRM-1 | 72.1 kD | 66.4 ± 0.3 kD | 14.46 |
| FRM-4 + FRM-1 | 136.7 kD | 68.7 ± 3.2 kD | 14.41 |

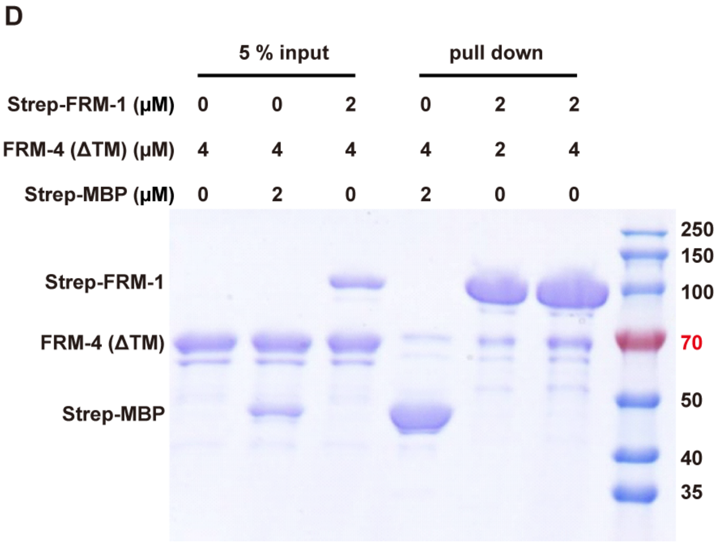


**Figure S5. Purified FRM-4 and FRM-1 exhibit both homophilic and heterophilic interaction *in vitro*, related to Figure 2**

(A and B) Size-exclusion chromatography coupled with multi-angle light scattering (SEC-MALS) analysis of purified FRM-1 (A) and FRM-4 (ΔTM) (B) at increasing protein concentrations. Both proteins exhibited concentration-dependent shifts toward smaller elution volumes, consistent with self-association.

(C) SEC-MALS analysis of a mixture of purified FRM-1 and FRM-4 (ΔTM), revealing an additional peak at a smaller elution volume than either protein alone, consistent with the formation of an FRM-1-FRM-4 complex.

(D) *In vitro* pull-down assay examining the interaction between FRM-1 and FRM-4. Purified Strep-tagged FRM-1 was used to pull down FRM-4 lacking its transmembrane domain. Strep-tagged maltose-binding protein served as a negative control. Input (5%) and pull-down fractions were analyzed by SDS-PAGE followed by Coomassie blue staining. Molecular weights (kDa) are indicated on the right.

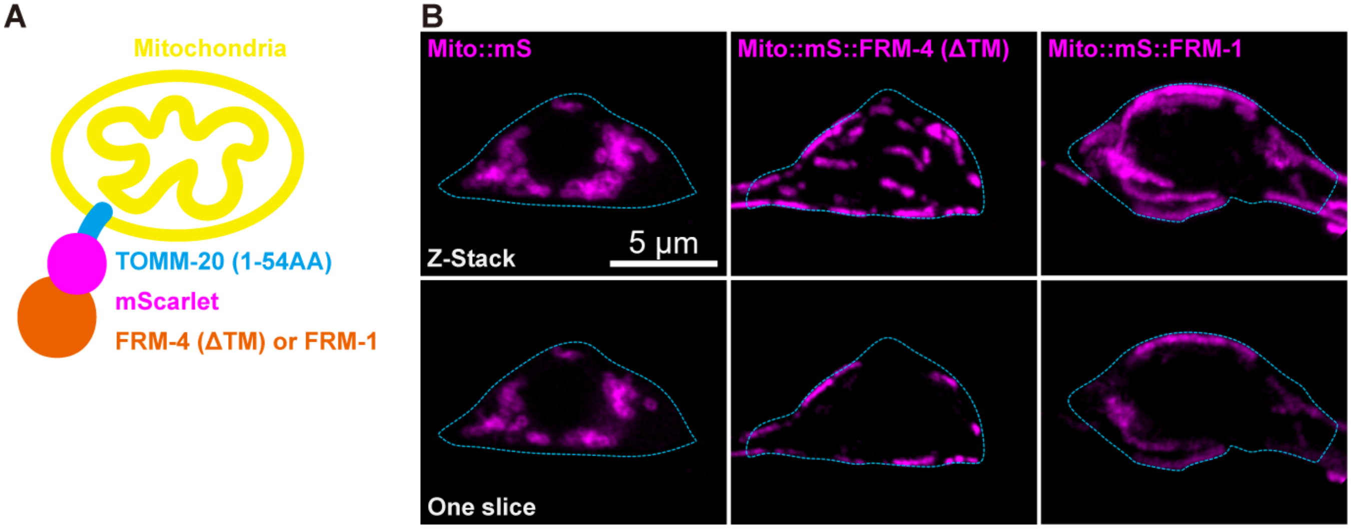


**Figure S6. FRM-4 and FRM-1 promote membrane contact formation, related to Figure 4**

(A) Schematics of mitochondrial targeting constructs comprising the N-terminal targeting sequence of TOMM-20 (amino acids 1-54) fused to FRM-4 (ΔTM) or FRM-1.

(B) Representative confocal z-stack projections and single optical sections showing mitochondrial localization in the PVD somas expressing TOMM-20 (1-54)::mScarlet alone or fused to FRM-4 (ΔTM) or FRM-1. All constructs were expressed from single-copy PVD-specific transgenes.

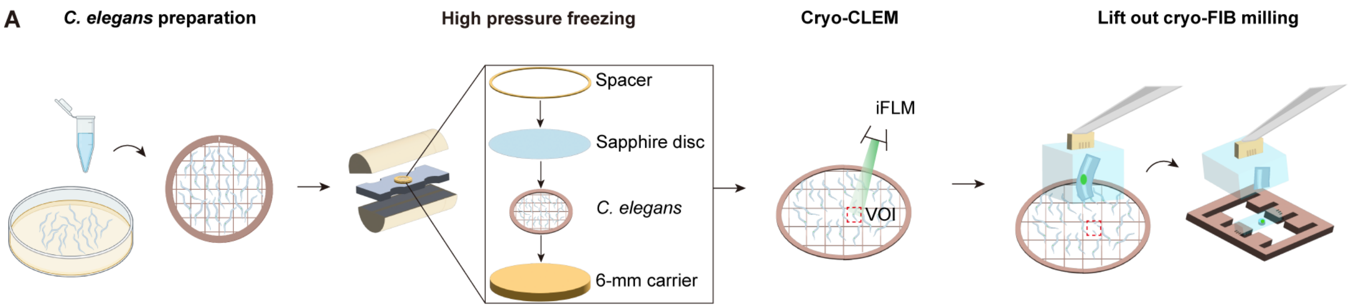


**Figure S7. Cryo-ET workflow for *C. elegans* sample preparation, related to Figure 5**

(A) Schematic of the cryo-ET workflow, including high-pressure freezing, cryo-CLEM localization of PVD neurons, cryo-FIB lift-out and milling, and cryo-ET data collection and reconstruction.

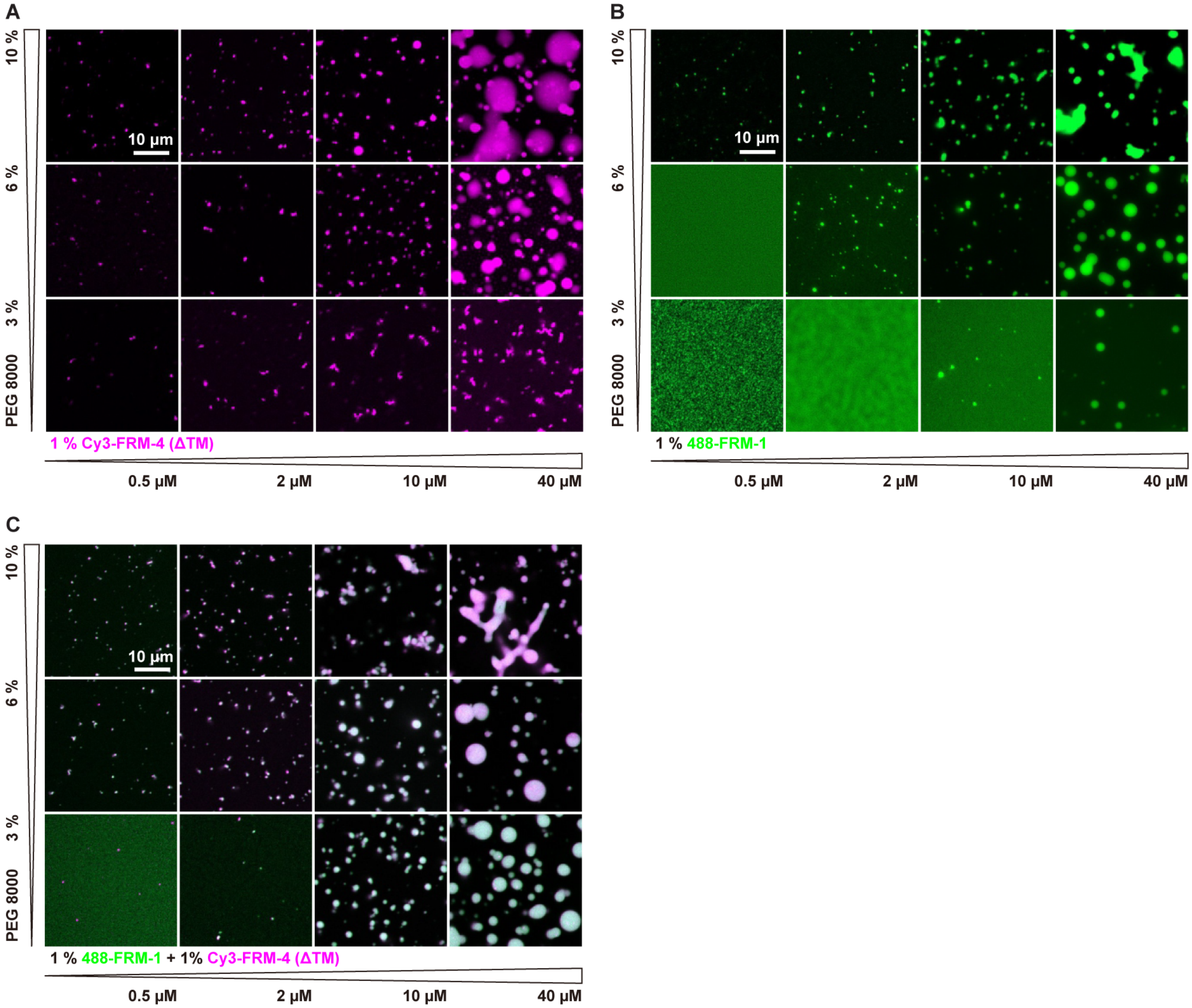


**Figure S8. Purified FRM-4 and FRM-1 undergo LLPS *in vitro*, related to Figure 6**

(A-C) Phase diagrams showing condensate formation by purified FRM-4 (ΔTM) (A), FRM-1 (B), and mixtures of FRM-1 and FRM-4 (ΔTM) (C) as a function of protein and PEG 8000 concentrations. Condensate formation was generally enhanced at higher protein and PEG concentrations. At elevated PEG concentrations, changes in condensate morphology were observed.

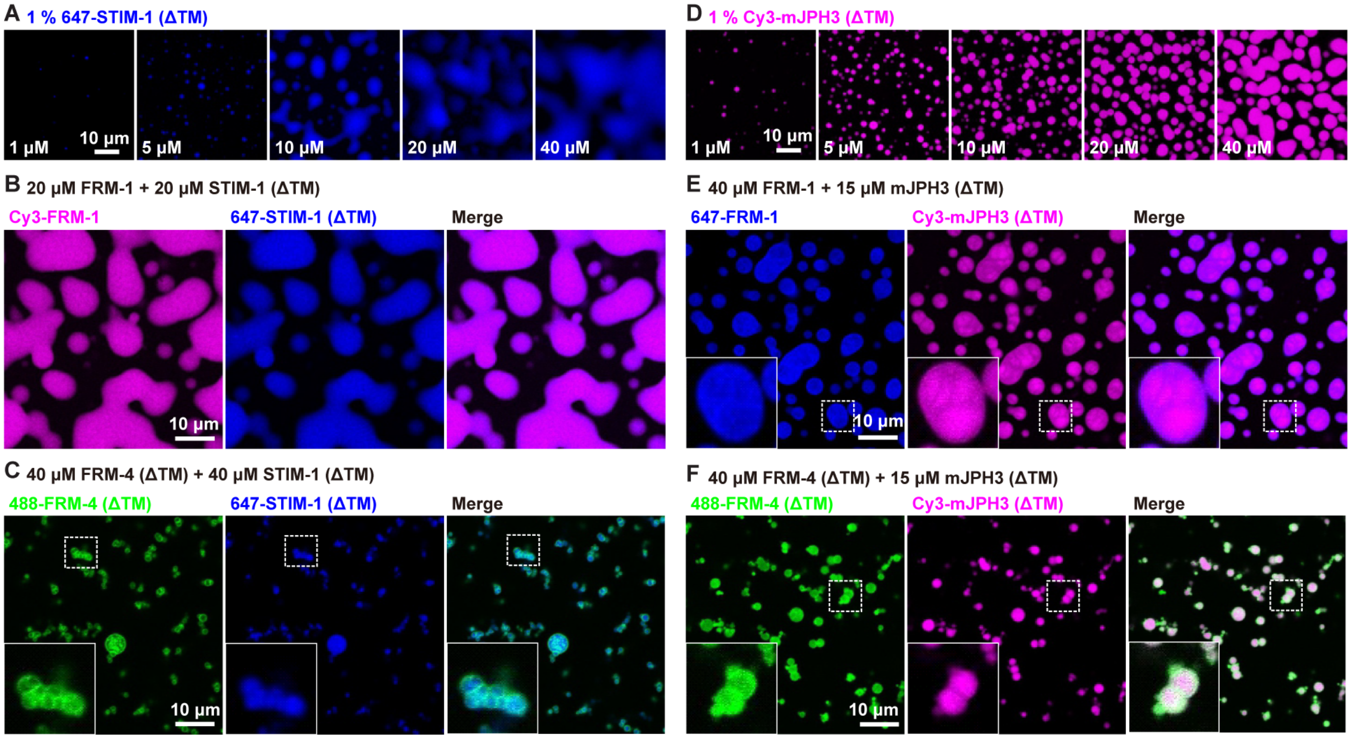


**Figure S9. Purified STIM-1 and mJPH3 exhibit selective mixing with FRM condensates *in vitro*, related to Figure 6**

(A) Phase diagram showing concentration-dependent condensate formation by purified STIM-1 (ΔTM).

(B) Representative confocal images showing condensates formed by mixtures of FRM-1 and STIM-1 (ΔTM) at the indicated protein concentrations.

(C) Representative confocal images showing the spatial segregation of STIM-1 (ΔTM) and FRM-4 (ΔTM) condensates at the indicated protein concentrations. The inset shows a threefold magnification of a representative condensate.

(D) Phase diagram showing concentration-dependent condensate formation by purified mJPH3 (ΔTM). The imaging buffer contained 20 mM HEPES (pH 7.5), 100 mM NaCl, and 1 mM TCEP (no extra PEG added).

(E and F) Representative confocal images showing the spatial segregation of mJPH3 (ΔTM) from condensates formed by FRM-1 (E) or FRM-4 (ΔTM) (F), at the indicated protein concentrations. Insets show threefold magnifications of representative condensates.

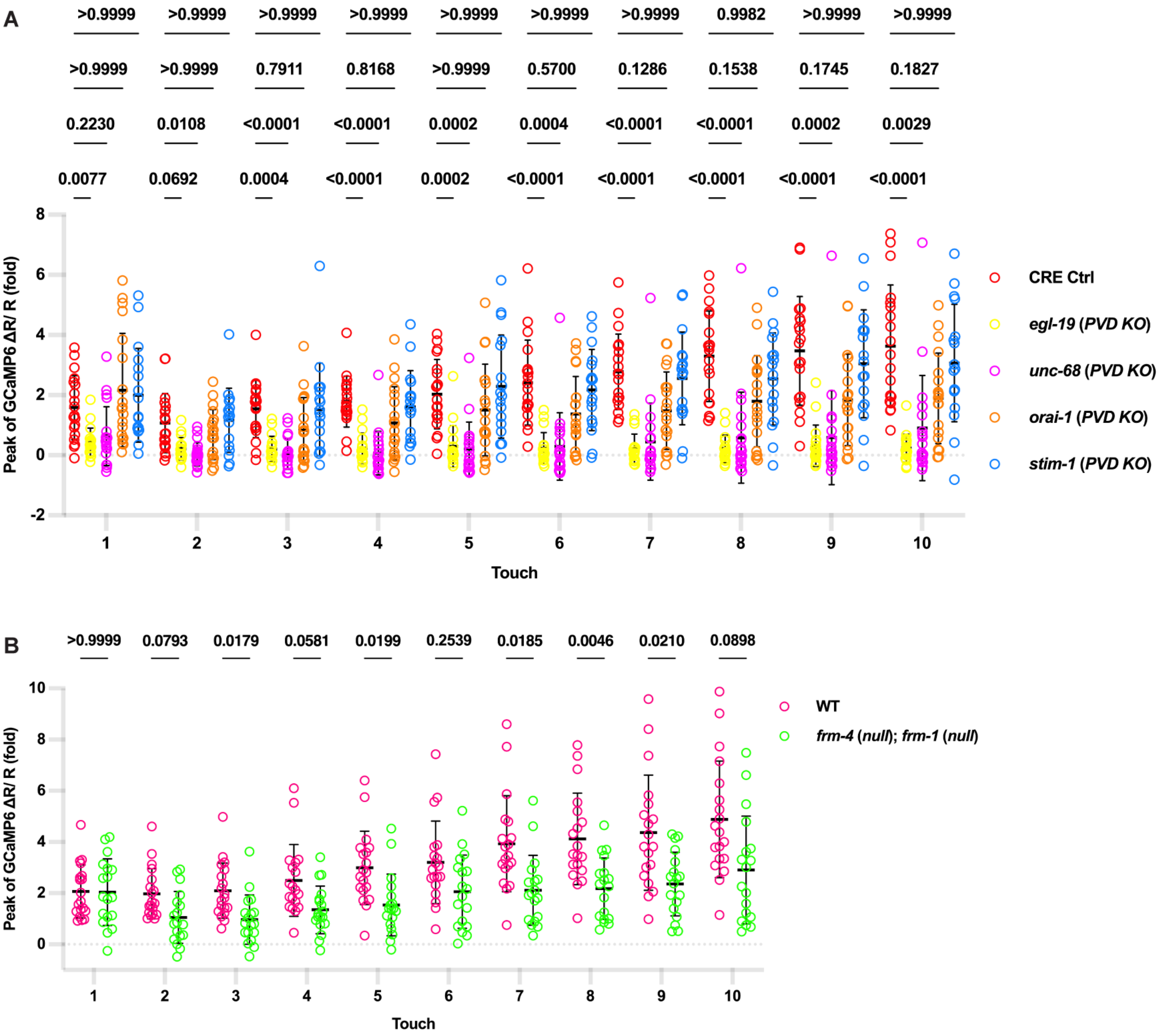


**Figure S10. Peak $Ca^{2+}$ responses to repeated harsh-touch stimulation, related to Figure 7**

(A) Quantification of peak GCaMP6 responses (ΔR/R) to each of 10 sequential harsh-touch stimuli in CRE control and PVD-specific *egl-19*, *unc-68*, *orai-1*, and *stim-1* knockout animals, corresponding to Figure 7A.

(B) Quantification of peak GCaMP6 responses (ΔR/R) to each of 10 sequential harsh-touch stimuli in WT and *frm-4*; *frm-1* animals, corresponding to Figure 7B.

For both panels, each dot represents one PVD neuron from an individual animal. Data are presented as mean ± SD. Statistical significance was determined using two-way repeated-measures ANOVA with Geisser-Greenhouse correction, followed by Šídák's multiple-comparisons test comparing each genotype at each stimulus.

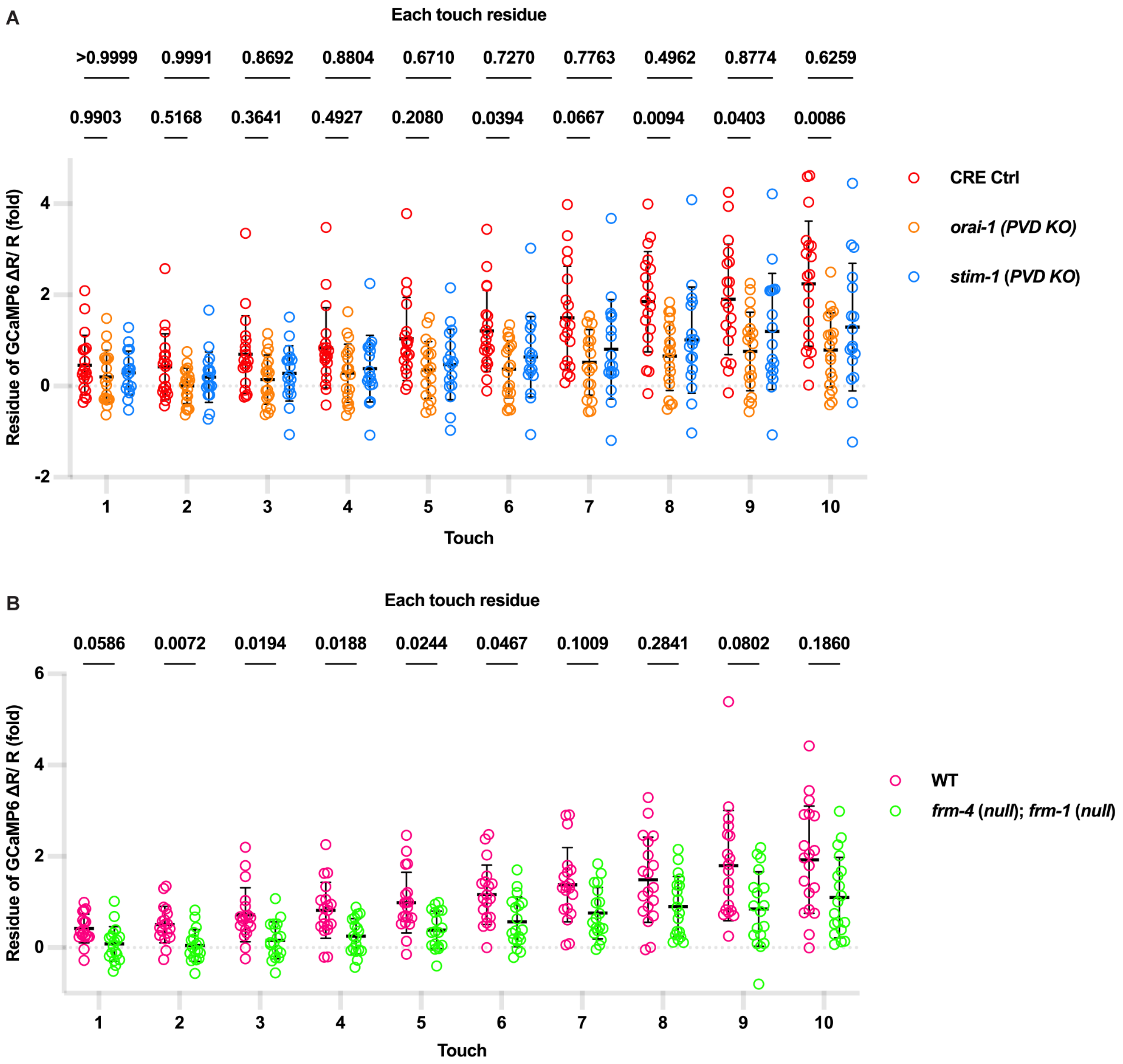


**Figure S11. Residual $Ca^{2+}$ signals following repeated harsh-touch stimulation, related to Figure 7**

(A) Quantification of residual GCaMP6 signals (ΔR/R), averaged over the 9-14 s interval following the onset of each of 10 sequential harsh-touch stimuli, in CRE control and PVD-specific *orai-1* and *stim-1* knockout animals, corresponding to Figure 7A.

(B) Quantification of residual GCaMP6 signals (ΔR/R) over the same interval in WT and *frm-4*; *frm-1* animals, corresponding to Figure 7B.

For both panels, each dot represents one PVD neuron from an individual animal. Data are presented as mean ± SD. Statistical significance was determined using two-way repeated-measures ANOVA with Geisser-Greenhouse correction, followed by Šídák's multiple-comparisons test comparing each genotype at each stimulus.

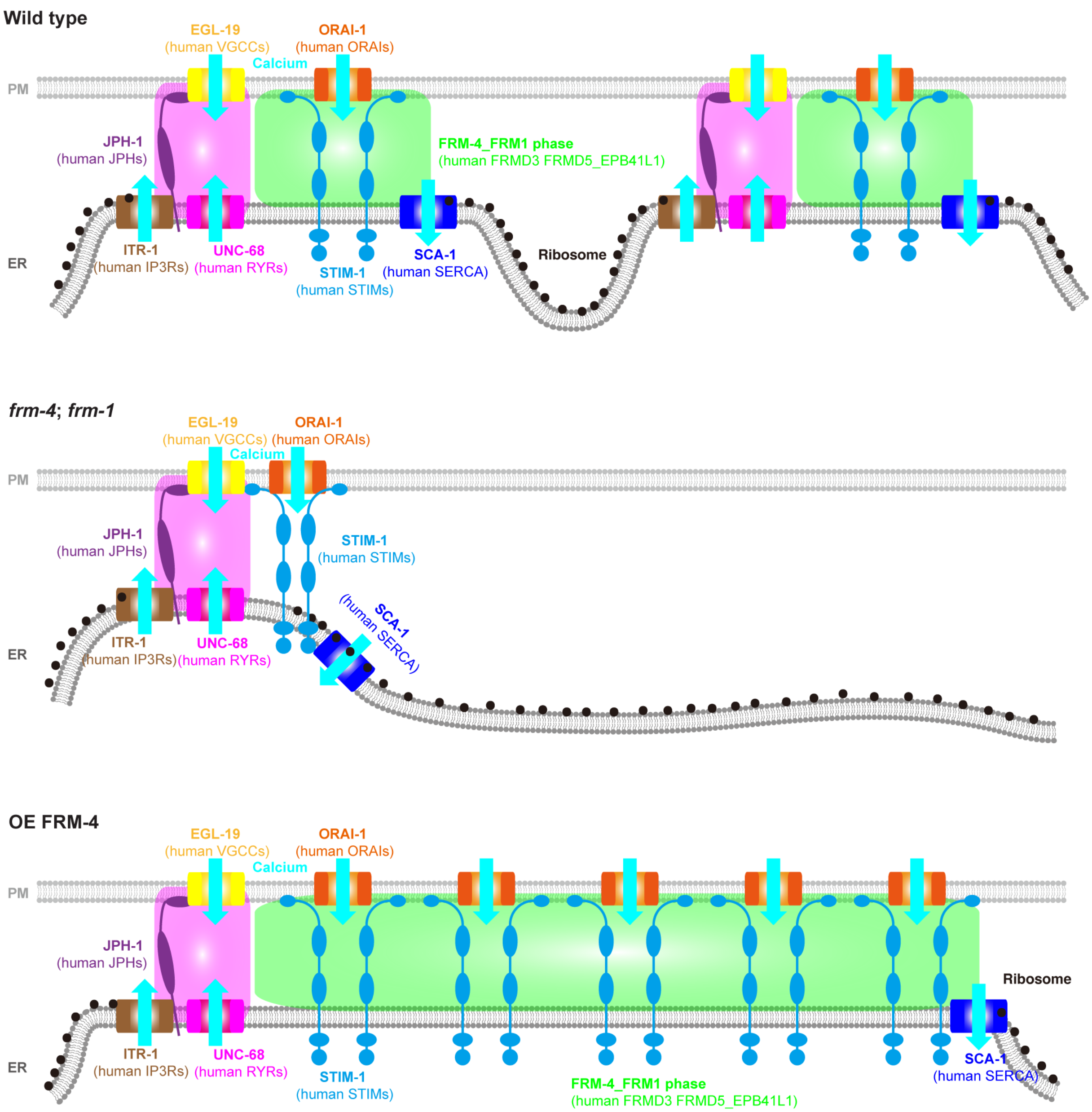


**Figure S12. Proposed model for FRM-dependent organization of $Ca^{2+}$-signaling subdomains at neuronal EPJs**

Schematic illustrating the organization of neuronal EPJs under wild-type conditions, following loss of FRM-4 and FRM-1, and upon FRM-4 overexpression.

Top, wild-type: FRM-4 and FRM-1 organize discrete EPJ patches containing adjacent, molecularly distinct $Ca^{2+}$-signaling subdomains. The SOCE subdomain contains ORAI-1 and STIM-1, whereas the CICR subdomain contains EGL-19, JPH-1, and UNC-68. Additional ER $Ca^{2+}$-handling proteins, including ITR-1 and SCA-1, are depicted.

Middle, *frm-4*; *frm-1* mutants: Loss of FRM-4 and FRM-1 disrupts EPJ architecture, reduces ER-PM proximity, and compromises the junctional organization of SOCE and CICR components, resulting in impaired neuronal $Ca^{2+}$ signaling.

Bottom, FRM-4 overexpression: Elevated FRM-4 expression expands EPJ-associated membrane contacts and redistributes $Ca^{2+}$-signaling components across an enlarged junctional domain.

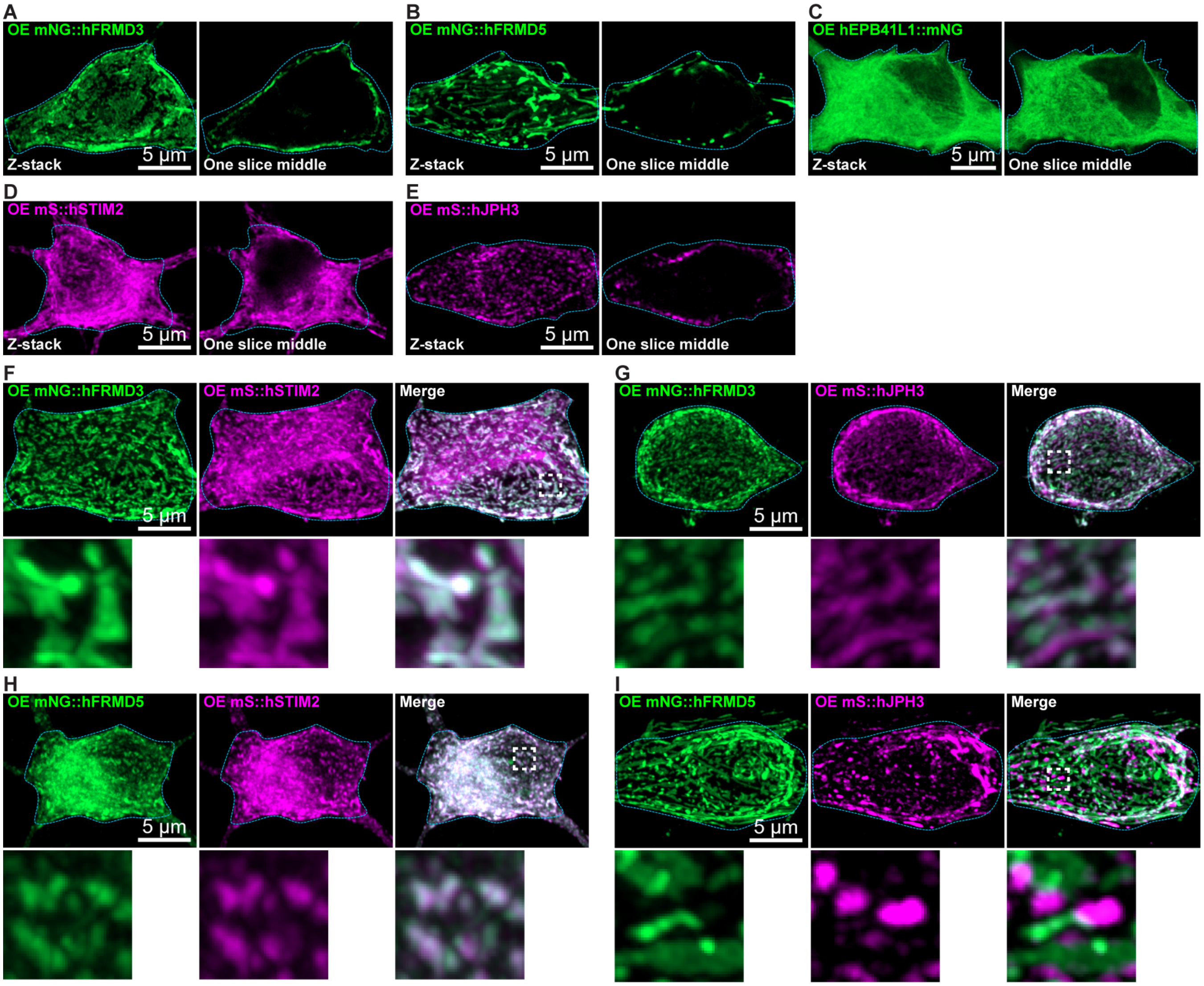


**Figure S13. FRMD3 and FRMD5 exhibit distinct spatial relationships with STIM2 and JPH3 in human neurons**

(A-C) Representative confocal z-stack projections and single optical sections showing the subcellular distributions of overexpressed mNeonGreen::hFRMD3 (A), mNeonGreen::hFRMD5 (B), and hEPB41L1::mNeonGreen (C) in the somas of human iPSC-derived neurons.

(D and E) Representative confocal z-stack projections and single optical sections showing the subcellular distributions of overexpressed mScarlet::hSTIM2 (D) and mScarlet::hJPH3 (E) in the somas of human iPSC-derived neurons.

(F and G) Representative confocal z-stack projections showing the spatial relationships between co-expressed mNeonGreen::hFRMD3 and mScarlet::hSTIM2 (F) or mScarlet::hJPH3 (G) in the somas of human iPSC-derived neurons.

(H and I) Representative confocal z-stack projections showing the spatial relationships between co-expressed mNeonGreen::hFRMD5 and mScarlet::hSTIM2 (H) or mScarlet::hJPH3 (I) in the somas of human iPSC-derived neurons.
