## supplemental table for "Phase-separated FRM proteins create signaling subdomains in inter-organelle junctions"

### Table S1. Strains, plasmids, crRNAs, and primers

#### Strains

| Figure / video | Strain name | Genotype | Notes | Citation |
|---|---|---|---|---|
| **Figure 1** | | | | |
| Figures 1A and 1B | N2 | | | |
| Figures 1F and 1G | TV32018 | orai-1 (wy2404) III; wySi1119 IV; wySi955 V; wySi927 V | orai-1 (wy2404) = orai-1::linker::mNG3Asp11X8. wySi1119 = [pdes-2_mNeonGreen3A(1-10)_unc-54 3'UTR]. wySi955 = [pdes-2::myr-BFP::let-858 3'UTR]. wySi927 = [AD2 cassette w/ punc-122::GFP]. | This paper |
| Figures 1F and 1G | TV32363 | stim-1 (wy2329) III; wySi1018 III; orai-1 (wy2404) III; wySi1119 IV; wySi955 V; wySi927 V | stim-1 (wy2329) = loxp::stim-1::loxp. wySi1018= [pnhr-81::CRE::tbb 3'UTR]. orai-1 stim-1 too close, so I re knock in loxp::stim-1::loxp in wySi1018 III; orai-1 (wy2404) III; wySi1119 IV; wySi955 V; wySi927 V. | This paper |
| Figures 1F and 1G | TV31762 | wySi1109 IV; wySi955 V; wySi927 V | wySi1109 = [ser-2Prom3::mStayGold::linker::stim-1 isoform A cDNA::unc-54 3'UTR]. | This paper |
| Figures 1F and 1G | TV31850 | orai-1 (wy2340) III; wySi1018 III; wySi1109 IV; wySi955 V; wySi927 V | orai-1 (wy2340) = lox2722::orai-1::lox2722. | This paper |
| Figures 1I and 1J | TV31352 | egl-19 (wy2300) IV; wySi955 V; wySi927 V | egl-19 (wy2300) = egl-19::linker::mStayGold. | This paper |
| Figures 1I and 1J | TV31384 | jph-1 (wy2298) I; wySi955 V; wySi927 V | jph-1 (wy2298) = mStayGold::linker::jph-1. | This paper |
| Figures 1I and 1J | TV31354 | unc-68 (wy2307) V; wySi955 V; wySi927 V | unc-68 (wy2307) = unc-68::linker::mStayGold. unc-68 wySi955 V; wySi927 V too close, so I re knock in unc-68::linker::mStayGold in wySi955 V; wySi927 V. | This paper |
| Figures 1I and 1J | TV31717 | egl-19 (wy2300) IV; frm-4 (wy2202) I; jph-1 (wy2337) I; wySi1018 III; wySi955 V; wySi927 V | frm-4 (wy2202) = mScarlet::linker::frm-4. jph-1 (wy2337) = loxp::jph-1::loxp. jph-1 frm-4 too close, so I re knock in mScarlet::linker::frm-4 in jph-1 (wy2337) I; wySi1018 III; wySi955 V; wySi927 V. | This paper |
| Figures 1I and 1J | TV31718 | unc-68 (wy2307) V; frm-4 (wy2202) I; jph-1 (wy2337) I; wySi1018 III; wySi955 V; wySi927 V | | This paper |
| Figure 1K | TV32119 | egl-19 (wy2300) IV; wySi1153 II; wySi955 V; wySi927 V | wySi1153 = [ser-2Prom3_orai-1 (C09F5.2a.1 cDNA)::linker::mScarlet I3::unc-54 3'UTR]. | This paper |
| **Figure S1** | | | | |
| Figure S1A | TV29118 | wySi935 V | wySi935 = [pdes-2::myr-GFP::let-858 3'UTR]. | This paper |
| **Figure S2** | | | | |
| Figure S2A | TV32018 | orai-1 (wy2404) III; wySi1119 IV; wySi955 V; wySi927 V | | This paper |

| Figure / video | Strain name | Genotype | Notes | Citation |
|---|---|---|---|---|
| Figure S2B | TV31762 | wySi1109 IV; wySi955 V; wySi927 V | | This paper |
| Figure S2C | TV32120 | wySi1109 IV; wySi1153 II; wySi955 V; wySi927 V | | This paper |
| Figure S2D | TV31352 | egl-19 (wy2300) IV; wySi955 V; wySi927 V | | This paper |
| Figure S2E | TV31384 | jph-1 (wy2298) I; wySi955 V; wySi927 V | | This paper |
| Figure S2F | TV31354 | unc-68 (wy2307) V; wySi955 V; wySi927 V | | This paper |
| Figure S2G | TV32337 | egl-19 (wy2300) IV; wySi1163 II; wySi955 V; wySi927 V | wySi1163 = [ser-2Prom3::mScarlet I3::linker::jph-1::unc-54 3'UTR]. | This paper |
| Figure S2H | TV32319 | unc-68 (wy2307) V; wySi1163 II; wySi955 V; wySi927 V | | This paper |
| **Figure 2** | | | | |
| Figure 2A | TV32017 | frm-4 (wy2396) I; wySi1125 I; wySi919 V | frm-4 (wy2396) = FRT (minimal) FLPon::mStayGold::frm-4. wySi1125 = [pdes-2::FLP::unc-54 3'UTR]. | This paper |
| Figure 2A | TV32221 | frm-1 (wy2021) I; frm-4 (wy2396) I; wySi1125 I; wySi919 V | frm-1 (wy2021) = frm-1 whole gene delete. | This paper |
| Figure 2A | TV32318 | frm-1 (wy2429) I; wySi1125 I; wySi919 V | frm-1 (wy2429) = frm-1::FRT (minimal) FLPon::mStayGold | This paper |
| Figure 2A | TV32365 | frm-4 (wy1998) I; frm-1 (wy2429) I; wySi1125 I; wySi919 V | frm-4 (wy1998) = frm-4 whole gene delete. | This paper |
| Figure 2C | TV32108 | wySi1108 II; frm-4 (wy2396) I; wySi1125 I; wySi955 V; wySi927 V | wySi1108 = [ser-2Prom3::mScarlet I3::MAPPER::unc-54 3'UTR]. | This paper |
| Figure 2D | TV32364 | wySi1108 II; frm-1 (wy2429) I; wySi1125 I; wySi955 V; wySi927 V | | This paper |
| **Figure S3** | | | | |
| Figure S3A | TV28625 | wyIs988 V | wyIs988 = [pDHC280 ser-2Prom3::GFP::MAPPER 1.5 ng/ul, pDHC270 ser-2Prom3::mScarlet 5 ng/ul, Pord-1::GFP 50ng/ul]. | This paper |
| Figure S3A | TV29114 | frm-4 (wy1931) I; wyIs988 | frm-4 (wy1931) = C24A11.8a.2:exon6:c.C472T:p.Q158X. | This paper |
| Figure S3A | TV29477 | frm-4 (wy1998) I; wyIs988 V | | This paper |
| Figure S3A | TV29516 | frm-4 (wy1998) I; wyIs988 V; wyEx10738 | wyEx10738 = [pDHC329 ser-2Prom3::frm-4 C24A11.8 DNA 1.5 ng/ul, myo-2::mCherry 2ng/ul, DNA Ladder]. | This paper |
| Figure S3A | TV29553 | frm-1 (wy2021) I; wyIs988 V | | This paper |
| Figure S3A | TV29539 | frm-2 (wy2017) III; wyIs988 V | frm-2 (wy2017) = frm-2 whole gene delete. | This paper |

| Figure / video | Strain name | Genotype | Notes | Citation |
|---|---|---|---|---|
| Figure S3A | TV30470 | erm-1(wy2084) I; wySi1018 III; wyIs988 V | erm-1(wy2084) = loxp::erm-1::loxp. | This paper |
| Figure S3A | TV29825 | frm-4 (wy1998) I; frm-1 (wy2021) I; wyIs988 V | | This paper |
| Figure S3A | TV29927 | frm-4 (wy1998) I; frm-2 (wy2017) III; wyIs988 V | | This paper |
| Figure S3A | TV30477 | frm-4 (wy1998) I; erm-1 (wy2084) I; wySi1018 III; wyIs988 V | | This paper |
| Figure S3D | TV32017 | frm-4 (wy2396) I; wySi1125 I; wySi919 V | | This paper |
| Figure S3E | TV32318 | frm-1 (wy2429) I; wySi1125 I; wySi919 V | | This paper |
| Figure S3G | TV32017 | frm-4 (wy2396) I; wySi1125 I; wySi919 V | | This paper |
| Figure S3G | TV32141 | frm-4 (wy2416) I; wySi1125 I; wySi919 V | frm-4 (wy2416) = FRT (minimal) FLPon::mStayGold::frm-4 (delete TM). | This paper |
| Figure S3I | TV32017 | frm-4 (wy2396) I; wySi1125 I; wySi919 V | | This paper |
| **Figure S4** | | | | |
| Figure S4D | TV31104 | frm-4 (wy2295) I | frm-4 (wy2295) = mStayGold::linker::frm-4. | This paper |
| Figure S4D | TV32317 | frm-4 (wy2396) I; wySi1125 I; wySi954 V | wySi954 = [prab-3::FLP::let-858 3'UTR]. | This paper |
| **Figure 3** | | | | |
| Figure 3A | TV32222 | orai-1 (wy2404) III; wySi1119 IV; frm-4 (wy2202) I; wySi955 V; wySi927 V | | This paper |
| Figure 3A | TV31815 | wySi1109 IV; frm-4 (wy2202) I; wySi955 V; wySi927 V | | This paper |
| Figure 3C | TV32018 | orai-1 (wy2404) III; wySi1119 IV; wySi955 V; wySi927 V | | This paper |
| Figure 3C | TV32147 | frm-4 (wy1998) I; frm-1 (wy2021) I; orai-1 (wy2404) III; wySi1119 IV; wySi955 V; wySi927 V | | This paper |
| Figure 3C | TV32366 | wySi1106 II; orai-1 (wy2404) III; wySi1119 IV; wySi955 V; wySi927 V | wySi1106 = [pdes-2::mScarlet I3::linker::frm-4::unc-54 3'UTR]. | This paper |
| Figure 3C | TV31762 | wySi1109 IV; wySi955 V; wySi927 V | | This paper |
| Figure 3C | TV31763 | frm-4 (wy1998) I; frm-1 (wy2021) I; wySi1109 IV; wySi955 V; wySi927 V | | This paper |
| Figure 3C | TV31828 | wySi1106 II; wySi1109 IV; wySi955 V; wySi927 V | | This paper |

| Figure / video | Strain name | Genotype | Notes | Citation |
|---|---|---|---|---|
| Figure 3E | TV31203 | frm-4 (wy2295) I; wySi919 V; unc-86 (wy1955) III | unc-86 (wy1955) = unc-86::P2A::mScarlet. | This paper |
| Figure 3E | TV31346 | orai-1 (wy2340) III; wySi1018 III; frm-4 (wy2295) I; wySi919 V | | This paper |
| Figure 3E | TV31441 | stim-1 (wy2329) III; wySi1018 III; frm-4 (wy2295) I; wySi919 V | | This paper |
| Figure 3E | TV31865 | orai-1 (wy2340) III; stim-1 (wy2329) III; wySi1018 III; frm-4 (wy2295) I; wySi919 V | orai-1 stim-1 too close, so I re knock in lox2722::orai-1::lox2722 in stim-1 (wy2329) III; wySi1018 III; frm-4 (wy2295) I; wySi919 V | This paper |
| Figure 3G | TV31885 | wySi1123 II; wySi955 V; wySi927 V | wySi1123 = [ser-2Prom3::tomm-20(1-54AA)::linker::mScarlet I3::unc-54 3'UTR]. | This paper |
| Figure 3G | TV31896 | wySi1126 II; wySi955 V; wySi927 V | wySi1126 = [ser-2Prom3::tomm-20(1-54AA)::linker::frm-4 (delete TM)::mScarlet I3::unc-54 3'UTR]. | This paper |
| Figure 3G | TV31897 | wySi1127 II; wySi955 V; wySi927 V | wySi1127 = [ser-2Prom3::tomm-20(1-54AA)::linker::frm-1::mScarlet I3::unc-54 3'UTR]. | This paper |
| **Figure 4** | | | | |
| Figure 4A | TV31757 | egl-19 (wy2300) IV; frm-4 (wy2202) I; wySi955 V; wySi927 V | | This paper |
| Figure 4A | TV31440 | jph-1 (wy2298) I; frm-4 (wy2202) I; wySi955 V; wySi927 V | | This paper |
| Figure 4A | TV31756 | unc-68 (wy2307) V; frm-4 (wy2202) I; wySi955 V; wySi927 V | | This paper |
| Figure 4C | TV31352 | egl-19 (wy2300) IV; wySi955 V; wySi927 V | | This paper |
| Figure 4C | TV31400 | frm-4 (wy1998) I; frm-1 (wy2021) I; egl-19 (wy2300) IV; wySi955 V; wySi927 V | | This paper |
| Figure 4C | TV31680 | wySi1106 II; egl-19 (wy2300) IV; wySi955 V; wySi927 V | | This paper |
| Figure 4C | TV31384 | jph-1 (wy2298) I; wySi955 V; wySi927 V | | This paper |
| Figure 4C | TV31401 | frm-4 (wy1998) I; frm-1 (wy2021) I; jph-1 (wy2298) I; wySi955 V; wySi927 V | jph-1 frm-4 too close, so I re knock in frm-4 whole gene delete in jph-1 (wy2298). | This paper |
| Figure 4C | TV31681 | wySi1106 II; jph-1 (wy2298) I; wySi955 V; wySi927 V | | This paper |
| Figure 4C | TV31354 | unc-68 (wy2307) V; wySi955 V; wySi927 V | | This paper |
| Figure 4C | TV31477 | frm-4 (wy1998) I; frm-1 (wy2021) I; unc-68 (wy2307) V; wySi955 V; wySi927 V | | This paper |

| Figure / video | Strain name | Genotype | Notes | Citation |
|---|---|---|---|---|
| Figure 4C | TV31682 | wySi1106 II; unc-68 (wy2307) V; wySi955 V; wySi927 V | | This paper |
| Figure 4E | TV31203 | frm-4 (wy2295) I; wySi919 V; unc-86 (wy1955) III | | This paper |
| Figure 4E | TV31339 | egl-19 (wy2338) IV; wySi1018 III; frm-4 (wy2295) I; wySi919 V | egl-19 (wy2338) = loxp::egl-19::loxp. | This paper |
| Figure 4E | TV31468 | jph-1 (wy2337) I;wySi1018 III; frm-4 (wy2295) I; wySi919 V | jph-1 (wy2337) = loxp::jph-1::loxp. frm-4 jph-1 too close, so I re knock in mStayGold::linker::frm-4 in loxp::jph-1::loxp. | This paper |
| Figure 4E | TV31355 | unc-68 (wy2341) V; wySi1018 III; frm-4 (wy2295) I; wySi919 V | unc-68 (wy2341) = lox2722::unc-68::lox2722. unc-68 wySi919 too close, so I re knock in lox2722::unc-68::lox2722 in wySi1018 III; frm-4 (wy2295) I; wySi919 V. | This paper |
| **Figure 5** | | | | |
| Figure 5A-5M | TV31333 | jph-1 (wy2298) I; wySi919 V | | This paper |
| Figure 5A-5M | TV31336 | frm-4 (wy1998) I; frm-1 (wy2021) I; jph-1 (wy2298) I; wySi919 V | | This paper |
| **Figure 6** | | | | |
| Figure 6A | TV31387 | frm-4 (wy2295) I; wySi955 V; wySi927 V | | This paper |
| Figure 6B | TV32018 | orai-1 (wy2404) III; wySi1119 IV; wySi955 V; wySi927 V | | This paper |
| **Figure 7** | | | | |
| Figure 7A, 7C, 7E, 7G | TV30474 | wySi1018 III; xuIs542 V | xuIs542 = [ser-2Prom3::GCaMP6(f), ser-2Prom3::mCherry]. | This paper |
| Figure 7A, 7C, 7E, 7G | TV32203 | egl-19 (wy2338) IV; wySi1018 III; xuIs542 V | | This paper |
| Figure 7A, 7C, 7E, 7G | TV32311 | unc-68 (wy2341) V; wySi1018 III; xuIs542 V | unc-68 xuIs542 too close, so I re knock in lox2722::unc-68::lox2722 in xuIs542. | This paper |
| Figure 7A, 7C, 7E, 7G | TV32158 | orai-1 (wy2340) III; wySi1018 III; xuIs542 V | | This paper |
| Figure 7A, 7C, 7E, 7G | TV32159 | stim-1 (wy2329) III; wySi1018 III; xuIs542 V | | This paper |
| Figure 7B, 7D, 7F, 7H | TQ8768 | xuIs542 V | | DOI: 10.1016/j.neuron.2021.08.035 |
| Figure 7B, 7D, 7F, 7H | TV30123 | frm-4 (wy1998) I; frm-1 (wy2021) I; xuIs542 V | | This paper |
| Video S1 | TV32018 | orai-1 (wy2404) III; wySi1119 IV; wySi955 V; wySi927 V | | This paper |

| Figure / video | Strain name | Genotype | Notes | Citation |
|---|---|---|---|---|
| Video S2 | TV32222 | orai-1 (wy2404) III; wySi1119 IV; frm-4 (wy2202) I; wySi955 V; wySi927 V | | This paper |
| Video S3 | TV31352 | egl-19 (wy2300) IV; wySi955 V; wySi927 V | | This paper |
| Video S3 | TV31384 | jph-1 (wy2298) I; wySi955 V; wySi927 V | | This paper |
| Video S3 | TV31354 | unc-68 (wy2307) V; wySi955 V; wySi927 V | | This paper |
| Video S4 | TV31440 | jph-1 (wy2298) I; frm-4 (wy2202) I; wySi955 V; wySi927 V | | This paper |

#### Table S1. Plasmids

| Plasmids | Citation |
|---|---|
| pDHC280 ser-2Prom3::GFP::MAPPER | This paper |
| pDHC270 ser-2Prom3::mScarlet | This paper |
| pDHC329 ser-2Prom3::frm-4 C24A11.8 DNA | This paper |
| pDHC494 pLV[Exp]-EF1A>mNeonGreen-3xGS-hFRMD3 | This paper |
| VB250219-1679ewh pLV[Exp]-EF1A>mNeonGreen-3xGS-hFRMD5 | This paper |
| pDHC496 pLV[Exp]-EF1A>hEPB41L1_3xGS_mNeonGreen | This paper |
| mScarlet-hSTIM2 | DOI: 10.1016/j.cell.2024.11.029 |
| pDHC529 pLV[Exp]-EF1A>mScarlet_3XGS_hJPH3 | This paper |

### Table S1. crRNAs and primers

| Mutants: | crRNA | | Primers | | | |
|---|---|---|---|---|---|---|
| orai-1 (wy2404) = orai-1::linker::mNG3Asp11X8. | CACTTGCTCCGTCAACAATTCGG | | TGTAAAGTTCACGGCAGTCG | CCTCTCTACAACTGGTGGGT | | |
| wySi1119 = [pdes-2_mNeonGreen3A(1-10)_unc-54 3'UTR]. | | | TCGCTACCATTACCAACTTGTC | GTTTACATTTATTGGGCCTGGATC | CCTTCCAATCCGCCATATCC | |
| stim-1 (wy2329) = loxp::stim-1::loxp. | TTTTCTTCAAATATATGACTTGG | CCACGTGGAATTTTCTCACTAGG | GAGATTCAGAGAAACAATCGGCAG | CAATAATCCACGAAACTCTACCCAT | CCGCCGAAATACATCGAAATCTAG | CAGTGCTCACGAGTAACCGT |
| wySi1018= [pnhr-81::CRE::tbb 3'UTR]. | | | ttctcgtctacttggtgcaac | GTCCCACAAGATCAACAAGTCA | catgcgtcgtatccgtaagg | |
| orai-1 (wy2340) = lox2722::orai-1::lox2722. | TAGATGTTTTTCTTGTTAAATGG | AAATGGCATGACGGAACACGTGG | TTTCCTAGCGTACTGGTGAC | GATGCTTAAAGGGCGATGTG | CTCTACTATGCCTACGGGTTC | GCTGAAACTGATTGCCACTT |
| egl-19 (wy2300) = egl-19::linker::mStayGold. | TTGTGGTGAGAAGAATCTTCTGG | | CGCTCGACGAGAAATTGAGG | GGAGAGGAGGCGAAGAGATTATT | | |
| jph-1 (wy2298) = mStayGold::linker::jph-1. | TGTGTCAGCAGTACGGCAAATGG | | actactccgtctatttccagtc | CATTGCCCTTGATAGGTGTTAC | | |
| unc-68 (wy2307) = unc-68::linker::mStayGold. | TGTGTCAGCAGTACGGCAAATGG | | GAGCACAACTTTGCCAATTACC | GCTAAGGCACGTTTGAAGTT | | |
| frm-4 (wy2202) = mScarlet::linker::frm-4. | ATCGGTTTTCGTGTTTCCAGGGG | | ATTCTAGCCTATACCTCCATTCTC | CGATGCTTATGGTGATCCTG | | |
| jph-1 (wy2337) = loxp::jph-1::loxp. | GGATTCAATGTGCAAGATGGCGG | CTGGTTTCACTTTTTTGAAATGG | GACTGTAGCTTCTCTGTGTTGC | GAAAGACGTAGTCGCTTGACT | TACTCAGACCCGAAATGTTACG | GAGCCTGTGACGAATGAGAT |
| frm-4 (wy2396) = FRT (minimal) FLPon::mStayGold::frm-4. | ATCGGTTTTCGTGTTTCCAGGGG | | ATTCTAGCCTATACCTCCATTCTC | CGATGCTTATGGTGATCCTG | | |
| wySi1125 = [pdes-2::FLP::unc-54 3'UTR]. | | | GTTCTGGAGTTCTACTCGTTGG | CAAGAGGAACACGACTTATCACA | TCGCTACCATTACCAACTTGTC | |
| frm-1 (wy2021) = frm-1 whole gene delete. | TGCAATAGAGAGTACTGATGCGG | ACGGAGGTTGAAAACAACTTTGG | TTCGGGCGATTGGAGAATAC | TTCAAACTGGCTACCCGATAC | CGAATTGAAGAGGACTTACCAGAAT | |
| frm-1 (wy2429) = frm-1::FRT (minimal) FLPon::mStayGold | TAATGATGAAAAGAATTATTGGG | | TGCCGGAATGTTTAGACTGAG | GTATCGGGTAGCCAGTTTGAA | | |
| frm-4 (wy1998) = frm-4 whole gene delete. | ATGGACTGATAACAACTGATAGG | TATTGCTGGTCGGAACCACGTGG | TCTTTGATGAGACAGTTCTCG | CCTTGTCGAAGTCGTAAGTC | CAGTGAATCCGCAAATCTTC | |

| **Mutants:** | **crRNA** | | **Primers** | | | |
|---|---|---|---|---|---|---|
| frm-4 (wy1931) = C24A11.8a.2:exon6:c.C472T:p.Q158X. | | | TCCAGCACGTCTAAGGGATC | GCTGGCCTTCTCCAGAAACT | | |
| frm-2 (wy2017) = frm-2 whole gene delete. | AAGATTGACAATTGGTTAGTTGG | AGTGCTAGAGTCTTCTAATCTGG | TTCTGGAAGATTCGTTCGACC | CGAAGAATGGACACAACTGATAATC | GAAGTCCAGTGAAGGGAAGGA | |
| erm-1 (wy2084) = loxp::erm-1::loxp. | CATGTGTCACTTTGAATTTTCGG | TTTTATTGATCATCTAGATCTGG | CCGGAAATAATAGCCCTGTACC | GTCCGAGTCGAAGATGGATTG | TTATTGTCCACCGGAAACCTC | GTCATAATACGGCGTTCCCA |
| frm-4 (wy2416) = FRT (minimal) FLPon::mStayGold::frm-4 (delete TM). | CAACGATGAGCAGAAGTATTAGG | | TGGATCTGGCGTTGATTTGG | CTTCCTAATACTTCTGCTCATCGTTG | CCAGCGAAGAGGATGAATTTGTATA | |
| frm-4 (wy2295) = mStayGold::linker::frm-4. | ATCGGTTTTCGTGTTTCCAGGGG | | ATTCTAGCCTATACCTCCATTCTC | CGATGCTTATGGTGATCCTG | | |
| wySi954 = [prab-3::FLP::let-858 3'UTR]. | | | AACCTTCGACCTCACTTTCC | gttagccaccttcttcacaca | CCAAGGAGGTCGAGAACTAAC | |
| egl-19 (wy2338) = loxp::egl-19::loxp. | GCTACCGTTTTTGACAACAATGG | GATATTGAGCATTTTTGAAATGG | ATATTGGCGGTTTGGCGAAG | GAATTGGAATCCAGGGAGGAG | AATCTCTTCGCCTCCTCTCC | GATACTGCGTAAAGGATGGGA |
| unc-68 (wy2341) = lox2722::unc-68::lox2722. | TTGGAGTTCTCAATCAAAGTTGG | GAATTCATGTGAACAAGAGAAGG | TTTCTTAGGTGTGTGTGACGTG | GGAGAGATGAAGGACGAGAAA | AATGTGTTACAGAAGAGGGTGG | GAAGCACGCCAATCTTGTTG |
